# A review of the relation between species traits and extinction risk

**DOI:** 10.1101/408096

**Authors:** Filipe Chichorro, Aino Juslén, Pedro Cardoso

**Affiliations:** LiBRE - Laboratory for Integrative Biodiversity Research, Finnish Museum of Natural History, University of Helsinki, PO Box 17 (Pohjoinen Rautatiekatu 13), 00014 Helsinki, Finland; Finnish Museum of Natural History, University of Helsinki, PO Box 17 (Pohjoinen Rautatiekatu 13), 00014 Helsinki, Finland

**Keywords:** biological traits, body size, habitat breadth, meta-analysis, geographical range, threat status

## Abstract

Biodiversity is shrinking rapidly, and despite our efforts only a small part of it has been assessed for extinction risk. Identifying the traits that make species vulnerable might help us to predict the outcome for those less known. We gathered information on the relations of traits to extinction risk from 173 publications, across all taxa, spatial scales and biogeographical regions, in what we think it is the most comprehensive compilation to date. Vertebrates and the Palaearctic are the most studied taxon and region because of higher accumulation of data in these groups. Among the many traits that have been suggested to be good predictors, our meta-analyses were successful in identifying two as potentially useful in assessing risk for the lesser-known species: regardless of the taxon, species with small range and habitat breadth are more vulnerable to extinction. On the other hand, body size (the most studied trait) did not present a consistently positive or negative response. In line with recent research, we hypothesize that the relationship between body size and extinction risk is shaped by different aspects, namely body size is a proxy for different phenomena depending on the taxonomic group.

## I. Introduction

The International Union for Conservation of Nature (IUCN) compiles and keeps updated a database with assessments of risk of extinction for species. As of January 2019, 26 840 (28%) of all 96 951 species in this list were either Critically Endangered, Endangered, or Vulnerable to extinction and 15 055 (16%) were Data Deficient (IUCN, 2019). Yet, species in the IUCN database mostly comprise well-known taxa (e.g. 67% of vertebrates have been assessed versus 0.8% of insects (IUCN, 2019)), and it will probably take decades until a reasonable proportion of many taxa, such as most invertebrates, are assessed (Cardoso *et al.*, 2011a, b, 2012). Increasing the number of species in the database to the point where we have an unbiased picture of extinction risk across all organisms during the next years seems highly unlikely, as is the Barometer of Life goal of assessing 160 000 species by 2020 (Stuart *et al.*, 2010). Moreover, extinction is taxonomically selective (e.g. 63% of cycads are assessed as threatened versus ‘only’ about 13% of bird species (IUCN, 2018)). The current proportions of endangered species might not represent the greater picture of species diversity. Therefore, alternative ways of predicting the risk of extinction of species are urgently needed.

Understanding which biological/ecological traits of species make them more vulnerable could help us predict their extinction risk and make species protection and conservation planning more efficient. This approach is not new. Some comparative studies can be traced back to the 19th century (see McKinney, 1997, for a thorough historical perspective), and since the beginning of the new millennium many new comparative studies have arisen on the topic, as well as discussions over their usefulness (Fisher & Owens, 2004; Cardillo & Meijaard, 2012; Murray *et al.*, 2014; Verde Arregoitia, 2016). Many traits have been tested across hundreds of publications. Body size, for example, was found to be positively correlated with extinction risk across multiple taxa (Seibold *et al.*, 2015; Terzopoulou *et al.*, 2015; Verde Arregoitia, 2016), either through direct effects (e.g. larger species require more resources) or as a proxy for other traits (e.g. larger species have slower life cycles and therefore respond more slowly to change). Range size and population density, even after considering that they are often used to quantify extinction risk, have also been extensively tested and found to be relevant, at least for mammals (Purvis et al., 2000a; González-Suárez, Gómez & Revilla, 2013; Bland *et al.*, 2015; Verde Arregoitia, 2016). Traits related to exposure to human pressures have also been relevant in predicting threats to species (Cardillo, 2003), and recently Murray *et al.* (2014) have called for more studies explicitly incorporating threats and the interplay between traits and threats into the analyses. The inclusion of threat information in predicting extinction risk has indeed proved to increase the explanatory power of models (Murray *et al.*, 2014), and in some cases the same trait can bolster extinction risk or prevent it, depending on the threat (González-Suárez *et al.*, 2013).

Most of the studies to date have focused on mammals (e.g. Purvis *et al.*, 2000a; Cardillo *et al.*, 2008; González-Suárez *et al.*, 2013) and other vertebrates (e.g. Owens & Bennett, 2000; Luiz *et al.*, 2016), with relatively few on plants (e.g. Sodhi *et al.*, 2008; Powney *et al.*, 2014; Stefanaki *et al.*, 2015) and invertebrates (e.g. Sullivan *et al.*, 2000; Koh, Sodhi & Brook, 2004; Arbetman *et al.*, 2017). Each study was made on different spatial settings and scales, testing different traits (often according to availability of data), and employed different methods and response variables. While this is necessary and valuable information, making sense of the plethora of contrasting results is difficult, and perceiving general trends and trying to cover current gaps and bias are urgent. In this work we attempt to answer the following questions through a comprehensive bibliography search, data exploration and meta-analysis:

- Which traits have been studied more often?
- Which traits have been suggested as predictors of extinction risk?
- How generalizable are the past results, i.e., are there traits that have a consistent response across taxonomical groups and geographical settings?

## II. Material and Methods

In this review we undertook two sequential analyses of studies that examined the relation between traits of species and their estimated extinction risk. The first one was an exploratory analysis of the literature, that allowed us to identify the traits have been more studied and which were found to be most relevant. The second analysis consisted of multiple meta-analyses, in which comparable data extracted from a subset of the studies were used to understand and quantify trends across studies and taxa from all published data and to see whether any general conclusions could be made from existing literature.

### (1) Bibliography selection

We were aiming to retrieve an extensive list of publications that explicitly performed comparative studies of biological/ecological traits and extinction risk/decline of species and to identify which traits, extrinsic factors and taxa were used in each analysis and at which spatial scale. In doing so, we first retrieved a list of candidate publications, and then we considered them or not for this review based on them meeting a set of criteria. To assemble the candidate list, we searched Web of Science using the keywords ‘trait*’ and ‘extinct*’ until July 2018, and we checked the abstracts and titles for appropriateness. Additionally, we collected all papers from previous similar reviews (Murray *et al.*, 2014; Verde Arregoitia, 2016), and included publications already known to us. To consider a given paper as relevant to our study, all the following conditions had to be met:

- more than five species were involved in the study;
- for each species there was information on at least one biological trait;
- for each species there was a measurement of its extinction risk;
- there was a statistical model linking the species traits (explanatory variables) to the extinction risk (response variables), assigning scores to each trait involved (not necessarily significance).

We considered as measurements of extinction risk:

- recent (anthropogenic) extinctions versus extant species;
- any variable (continuous, ordinal, categorical or binary) directly indicating relative extinction risk, whether it was based on the IUCN Red List categories or not;
- population trend data, or a proxy of population trend data, in time;
- any other variables that indicated decline of species over time and/or risk of extinction.

### (2) Data collection

We assembled information on each comparative statistical test employed in each article. For each of these tests, we extracted information on the following (see also Table 1 and Appendix S1, S2):

- **Taxonomic group**: mammals, birds, reptiles, amphibians, fishes, insects, molluscs, other invertebrates, plants and fungi.
- **Geographical realm**: Afrotropics, Antarctic, Australasia, Indo-Malaya, Nearctic, Neotropics, Oceania and Palaearctic (Olson *et al.*, 2001).
- **Traits**: continuous, ordinal, categorical or binary – units, the number of observations (usually species), and whether there was a significant response to extinction risk for that test. We grouped traits with similar biological meaning into the same unified trait (e.g. body length and body mass into ‘body size’, all original names and assigned unified traits are available in Appendix S2). Henceforth, trait refers to these unified traits.

**Table 1:**
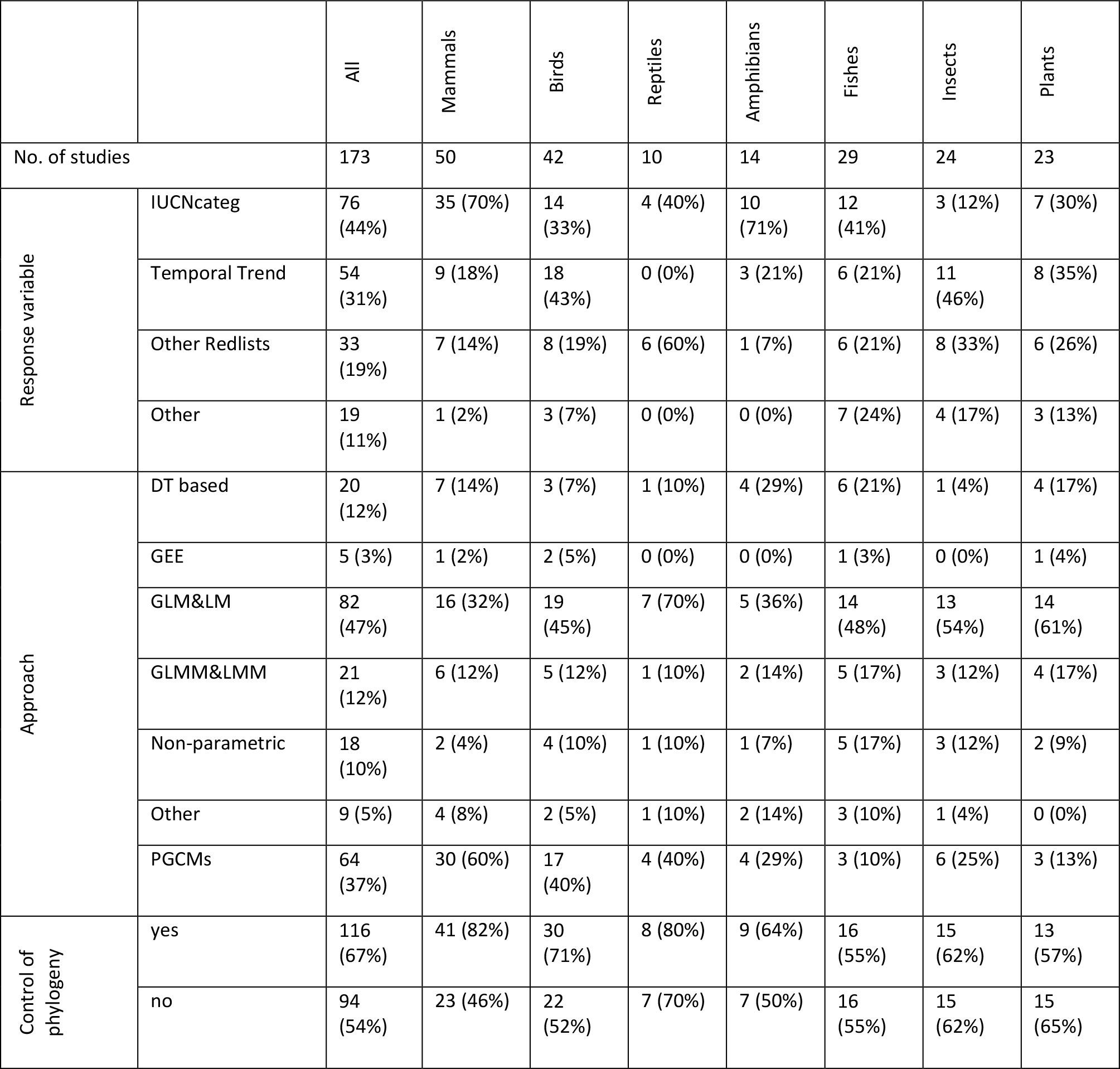
Number of publications within each taxonomic group, response variables, statistical approach, and controlling phylogeny or not. Percentages are of the number of publications within each category divided by the total number of publications in the given taxonomic group. GEE = generalized estimating equation; GLM = generalized linear model; GLMM = generalized linear mixed model; LM = linear model; LMM = linear mixed model; PGCM = phylogenetic comparative method. **Control of phylogeny**: we distinguished absolute no control of phylogeny (no) from at least some control of phylogeny (yes, via using phylogenetic trees or a taxonomical higher group as a controlling or covariable in the analyses).

|  |  | All | Mammals | Birds | Reptiles | Amphibians | Fishes | Insects | Plants |
| --- | --- | --- | --- | --- | --- | --- | --- | --- | --- |
| No. of studies |  | 173 | 50 | 42 | 10 | 14 | 29 | 24 | 23 |
| Response variable | IUCNcateg | 76 (44%) | 35 (70%) | 14 (33%) | 4 (40%) | 10 (71%) | 12 (41%) | 3 (12%) | 7 (30%) |
|  | Temporal Trend | 54 (31%) | 9 (18%) | 18 (43%) | 0 (0%) | 3 (21%) | 6 (21%) | 11 (46%) | 8 (35%) |
|  | Other Redlists | 33 (19%) | 7 (14%) | 8 (19%) | 6 (60%) | 1 (7%) | 6 (21%) | 8 (33%) | 6 (26%) |
|  | Other | 19 (11%) | 1 (2%) | 3 (7%) | 0 (0%) | 0 (0%) | 7 (24%) | 4 (17%) | 3 (13%) |
| Approach | DT based | 20 (12%) | 7 (14%) | 3 (7%) | 1 (10%) | 4 (29%) | 6 (21%) | 1 (4%) | 4 (17%) |
|  | GEE | 5 (3%) | 1 (2%) | 2 (5%) | 0 (0%) | 0 (0%) | 1 (3%) | 0 (0%) | 1 (4%) |
|  | GLM&LM | 82 (47%) | 16 (32%) | 19 (45%) | 7 (70%) | 5 (36%) | 14 (48%) | 13 (54%) | 14 (61%) |
|  | GLMM&LMM | 21 (12%) | 6 (12%) | 5 (12%) | 1 (10%) | 2 (14%) | 5 (17%) | 3 (12%) | 4 (17%) |
|  | Non-parametric | 18 (10%) | 2 (4%) | 4 (10%) | 1 (10%) | 1 (7%) | 5 (17%) | 3 (12%) | 2 (9%) |
|  | Other | 9 (5%) | 4 (8%) | 2 (5%) | 1 (10%) | 2 (14%) | 3 (10%) | 1 (4%) | 0 (0%) |
|  | PGCMs | 64 (37%) | 30 (60%) | 17 (40%) | 4 (40%) | 4 (29%) | 3 (10%) | 6 (25%) | 3 (13%) |
| Control of phylogeny | yes | 116 (67%) | 41 (82%) | 30 (71%) | 8 (80%) | 9 (64%) | 16 (55%) | 15 (62%) | 13 (57%) |
|  | no | 94 (54%) | 23 (46%) | 22 (52%) | 7 (70%) | 7 (50%) | 16 (55%) | 15 (62%) | 15 (65%) |

### (3) Exploratory analysis

In the exploratory analysis we first compared the number of studies across taxa, biogeographical realms, proxy of extinction used, statistical methodology, and if phylogeny was controlled for. Next, we compared the number of studies and the number of measurements (the number of measurements corresponds to the total number of statistical coefficients of each trait, usually corresponding to the number of statistical tests for that trait) in which each trait was used and calculated the percentage of significant measurements of each trait. Statistical tests which did not assign significance levels to traits had to be excluded from this step (e.g. most decision tree methodologies).

### (4) Meta-analysis

To understand whether traits were positively or negatively related to extinction risk across the multiple studies, we performed meta-analyses for each continuous trait. Meta-analyses are useful because they allow the comparison of outcomes from different studies by converting the outcomes to effect sizes. The use of Fisher’s Z as the effect size has the advantage of allowing very diverse statistical methodologies into the same effect size measurement. Effect sizes were obtained by transforming the statistics reported in the manuscripts (F, z, X^2^, t or r^2^) into Pearson’s product-moment correlation coefficients (r) by applying equations (1) to (5) (Rosenthal, 1991) and then transforming r into Fisher’s Z using equation (6) using R package metafor (Viechtbauer, 2010):

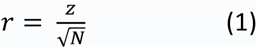

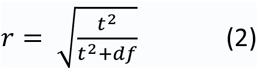

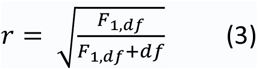

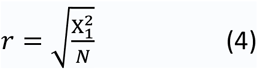

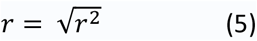

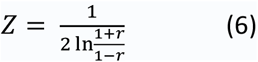

To ensure that the outcomes would be comparable, we restricted the analyses to univariate tests. To detect the overall effect size for each trait, we run linear mixed models. In relation to more traditional analytic tools, mixed models can be more flexible in controlling multiple measurements within studies (and hence non-independence of observations) through the use of random effects (see Prugh, 2009; Chaplin-Kramer *et al.*, 2011). Fisher’s Z was the response variable and was weighted by the inverse of the sample sizes. The response variable was tested against the intercept term only, with random effects being taxonomic group and study.

## III. Results

A total of 173 manuscripts fulfilled all criteria and were included in this study (Appendix S1).

### (1) Exploratory analysis

#### (a) Studies

The number of publications relating traits to extinction risk has increased steadily (Fig. 1). Mammals and birds have received the most attention over the years, followed by fishes, insects and plants (Table 1). Most studies were conducted in the Palaearctic region (Fig. 2), particularly for insects. Of particular note, amphibians, reptiles and mammals have been included in many studies focusing on the Australasian realm. Oceania and especially the Antarctic were the least represented biogeographical realms, with intermediate values in all the other regions.

**Figure 1:**
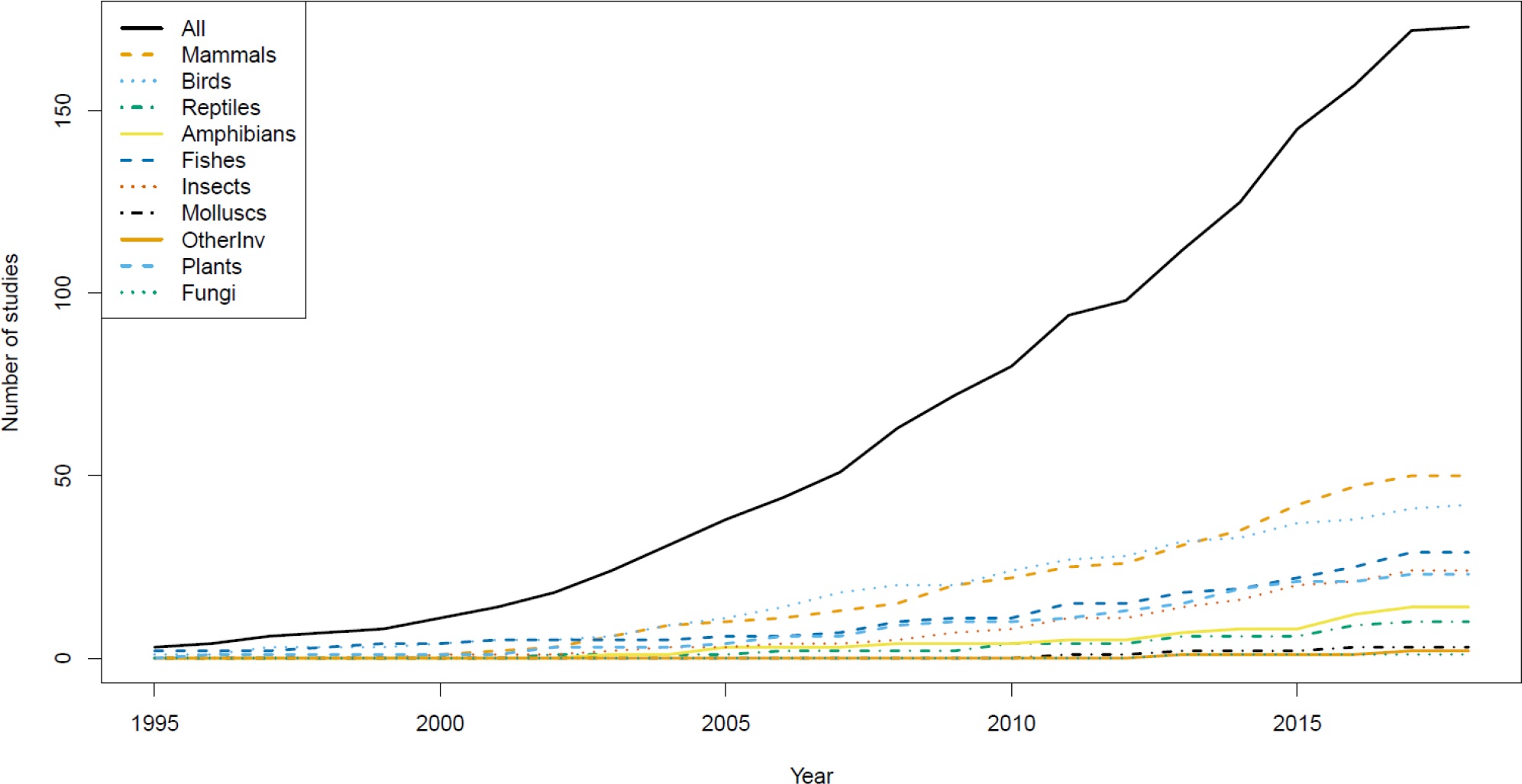
Cumulative number of publications per taxon relating traits to extinction risk.

**Figure 2:**
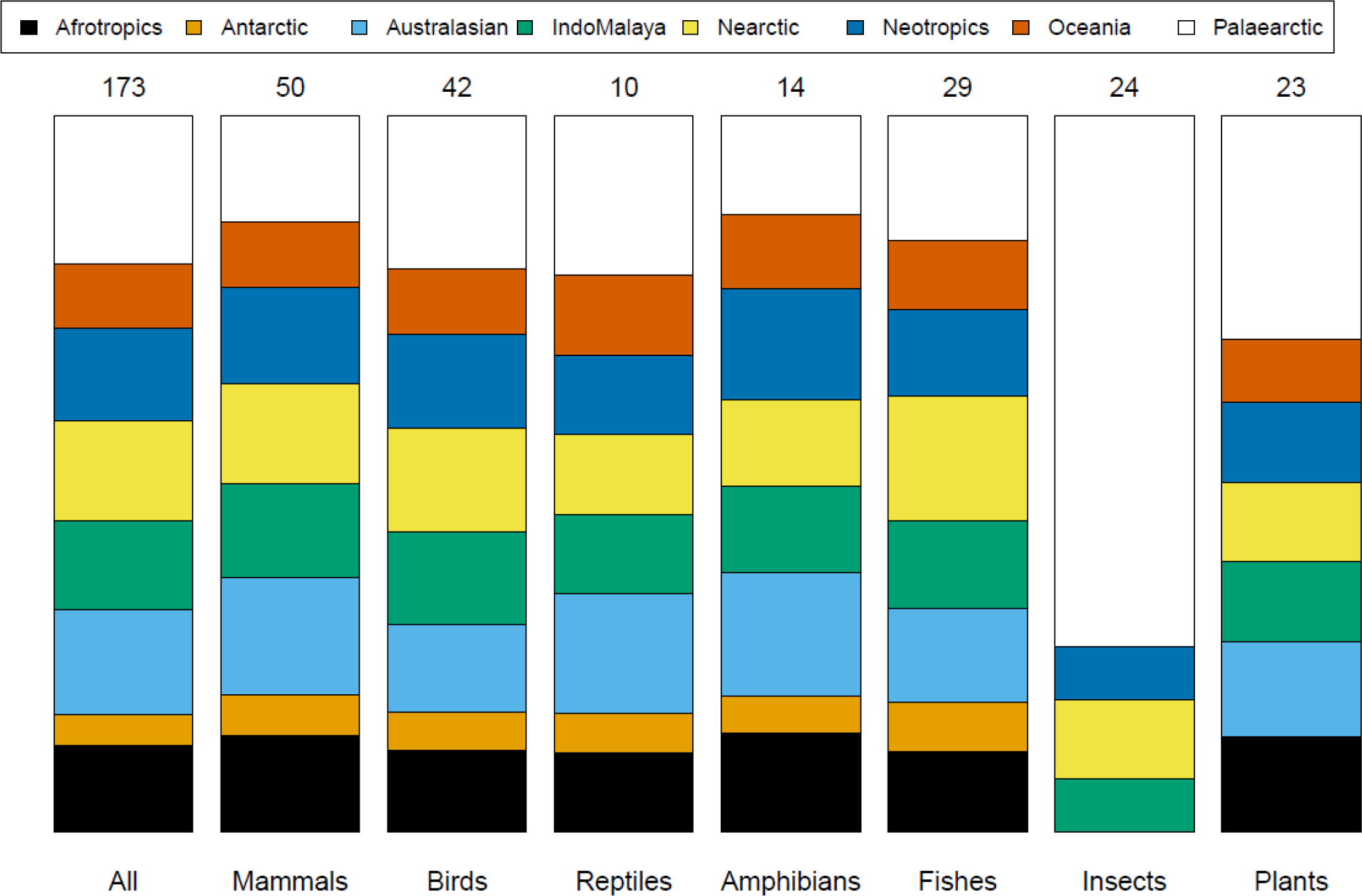
Proportion of manuscripts focusing on the different biogeographical regions by taxonomic group. Numbers above columns are the total number of studies per taxon.

#### (b) Traits

Body size was by far the most studied trait (Fig. 3, Table S1), followed by geographical range size and fecundity. Among the traits that were present in at least 10% of the studies, geographical range size was the trait with the greatest proportion of studies with significant measurements (almost three quarters) (Fig. 3). Besides geographical range size, only location (the geographical setting of the study) was significant in at least half of the measurements, but many traits were significant in >40% of the tests: body size, habitat type, diet breadth, habitat breadth, temperature and microhabitat type (Fig. 3). Fecundity, while amongst the most tested traits, was significant in only 27% of the measurements.

**Figure 3:**
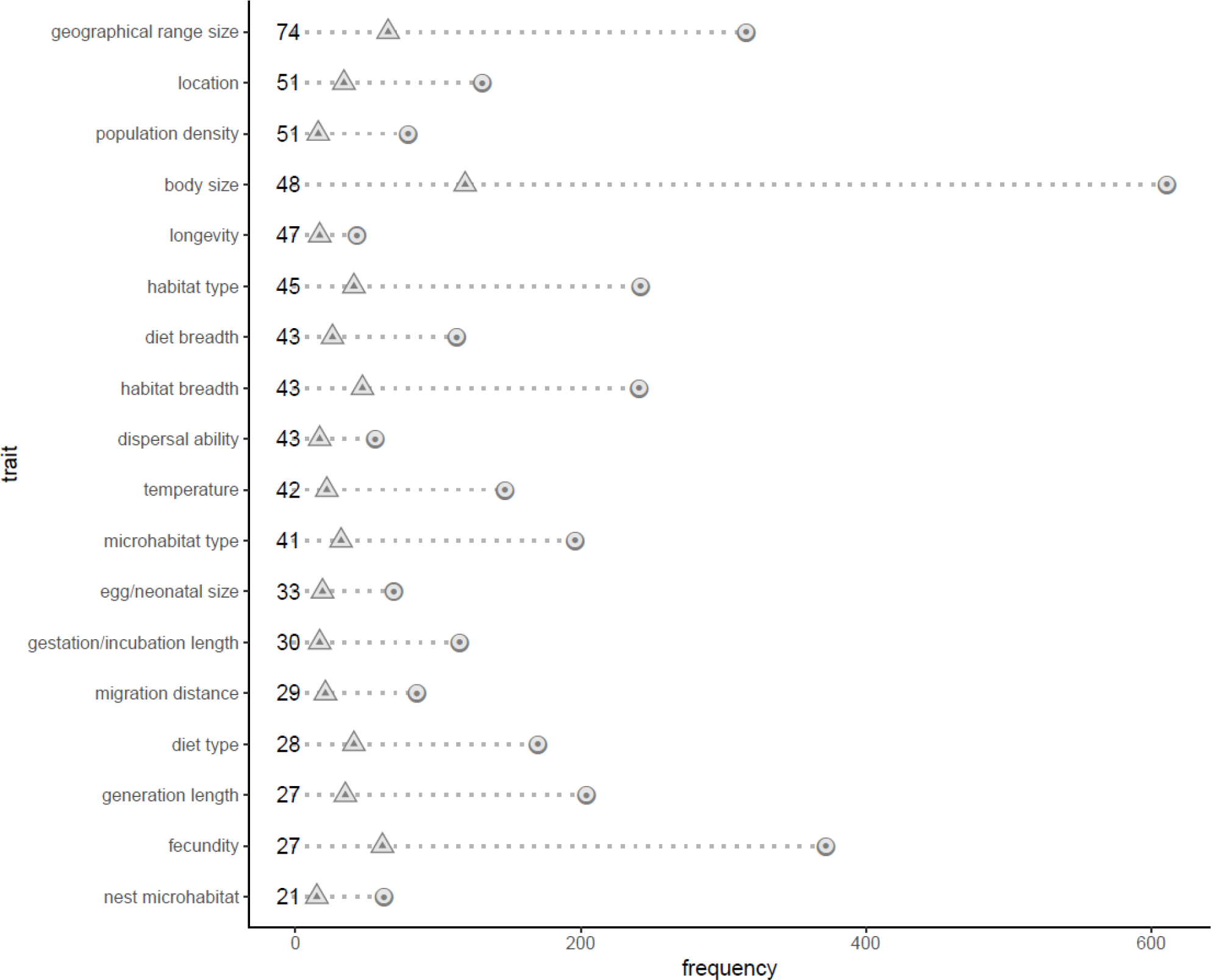
Summary information on variable use among all studies, depicting only variables included in at least 17 (10%) studies. The numbers before the dotted lines indicate the percentage of measurements in which the variable was significant. **Triangles**: number of studies in which the variable appears. **Circles**: total number of measurements for that variable.

Even when used in at least 10% of studies, not all of these traits were studied across all taxa. Body size and geographical range size were the only traits that were studied for all taxa (except for fungi, since the only study focusing on fungi did not attribute significance levels to traits and thus this group was not included here) and were significant in at least one test for each taxon (Appendix S3, Fig. S1 – S4).

Despite occurring in less than 10% of the studies, many traits have been found to be good predictors of extinction risk for some taxa. A number of traits (see Appendix S3 for the significances of all tested traits within taxa) were tested in at least three studies and were significant at least once, even if for single taxa (torpor/hibernation and weaning age in mammals; duration of flight period in birds; temperature for breeding in fishes; overwintering stage in insects; pollen vector, reproduction type, dispersal agent, and seed size in plants).

### (2) Meta-analysis

Geographical range size, habitat breadth, and body size were the only traits from which we could determine effect sizes and sample sizes from at least 10 studies including univariate tests – the minimum number that we considered reasonable in order to have confidence in the results of the meta-analyses. Effect sizes of geographical range size and body size mostly originated from mammal and bird studies but also from studies on reptiles, amphibians, fishes, insects, other invertebrates and plants (Appendix S4, Figs S5, S6). Effect sizes of habitat breadth also originated mostly from mammal and bird studies, yet reptile, amphibian, other invertebrates and plant studies were included (Appendix S4, Fig. S7).

For geographical range size and habitat breadth, the overall effect size was consistently and significantly negative across taxa and studies (Table 2, Appendix S4, Figs S5, S7). Contrastingly, the linear mixed model revealed an overall effect size not different from zero for body size (Table 2). Effect sizes of body size were either positive or negative (Appendix S4, Fig. S6), and while there was some tendency in mammals and birds for the effect sizes to be positive, although not consistently so, the effect sizes for plants and other invertebrates were strongly negative.

**Table 2:** Results of the linear mixed-effect models relating extinction risk with body size, geographical range size and habitat breadth. **df:** Degrees of freedom, **P:** p value.

| Model | Number of studies /<br>measurements /<br>different taxa | Taxa (number of<br>studies/measurements) | Intercept<br>estimate<br>(standard<br>error) | *df* | t value | *P* |
| --- | --- | --- | --- | --- | --- | --- |
| Effect sizes of Geographical range size ~<br>(Intercept) + Random(Study) +<br>Random(Taxon) | 23/49/9 | Mammals (8/12), Birds<br>(7/17), Reptiles (3/4),<br>Amphibians (3/4), Fishes<br>(2/4), Vertebrates (1/2),<br>Insects (2/4), Other<br>Invertebrates (1/1),<br>Plants (1/1) | -0.480 (0.107) | 24.787 | -4.479 | 0.0001 |
| Effect sizes of body size ~ (Intercept) +<br>Random(Study) + Random(Taxon) | 31/85/9 | Mammals (9/33), Birds<br>(11/26), Reptiles (4/9),<br>Amphibians (3/3), Fishes<br>(3/7), Vertebrates (1/1),<br>Insects (1/1), Other<br>invertebrates (1/1),<br>Plants (3/4) | 0.130 (0.104) | 11.000 | 1.251 | 0.237 |
| Effect sizes of Habitat breadth ~<br>(Intercept) + Random(Study) +<br>Random(Taxon) | 14/27/6 | Mammals (2/6), Birds<br>(6/13), Reptiles (2/3),<br>Amphibians (2/3),<br>Vertebrates (1/1), Other<br>invertebrates (1/1),<br>Plants (1/1) | -0.210 (0.041) | 11.843 | -5.112 | 0.0003 |

## IV. Discussion

Our review clearly reveals the increasing importance of the study of species traits on the understanding and prediction of extinction risk. The interest in the subject, even if relatively recent, is increasing exponentially and shows no signs of slowing down. Yet, we also found that past studies were biased in scope in terms of taxa, with vertebrates having the largest share, and spatial setting, with the Palaearctic dominating across taxa, although Australasia is much studied for mammals, reptiles and amphibians. Such biases should be mostly due to a large body of accumulated knowledge on these taxa and regions, to which a predominance of researchers in these areas continue to contribute. The special interest in the Australasian mammals may be due to an ongoing debate on the role of body size in extinction risk in this particular region (Verde Arregoitia, 2016).

Despite being, to our knowledge, the largest review of the relation between traits and extinction risk to date, we are aware that this contribution might not include all relevant studies in this field. The proportion of non-vertebrate studies included in our study is larger than that of Murray *et al.* (2014), even though the bias is inevitable in any comprehensive study on this subject. We are, however, confident that this review is thorough and as unbiased as possible with current data.

### (1) Relation between traits and extinction risk

Geographical range size was the best predictor of extinction risk overall. This is not surprising, since small geographical range is one of the criteria used in IUCN assessments (criteria B, D2), and these assessments are the measure of extinction risk in many studies, which might lead to circular reasoning. However, even when excluding from the analyses all species considered threatened due to small range, range was still strongly associated with extinction risk (e.g. Purvis *et al.*, 2000a; Wang *et al.*, 2018). The mechanism behind this relationship is not entirely understood (Purvis *et al.*, 2000a), but geographical range size captures ecological and dispersal attributes of species that would require harder to obtain variables, such as overall abundance of species, which are important in understanding extinction risk (Polaina, Revilla & González-Suárez, 2016). The abundance–occupancy relationship is a well-known and thoroughly studied pattern, and many mechanisms relate abundance to extinction risk (Gaston *et al.*, 2002). Likewise, range size is related to the dispersal ability of species, determining the capacity of a species to occupy new areas to escape multiple pressures, and with habitat breadth, revealing the ability of a species to cope with habitat change or loss.

Among the studies we included in our analysis, species with greater habitat breadth (habitat generalists) were less prone to becoming extinct. Specialists have long been regarded as losers, and generalists as winners in the current extinction crisis (McKinney & Lockwood, 1999; Clavel, Juliard & Devictor, 2011). Whether this trend is due to the intrinsic specificity of the species or to geographical range size is, however, not trivial to discern. In the studies included in this review, most habitat breadth measures were derived from maps. Consequently, less widespread species have less sampling points and therefore might show smaller habitat breadth due to sampling bias alone (Burgman, 1989), when in reality we lack knowledge of whether they could thrive under different habitats. Nonetheless, Slatyer, Hirst & Sexton (2013) showed that even after taking into consideration sampling bias, the relationship between habitat breadth and geographical range size remains significant across taxa. Irrespective of the putative causes or relations to other variables, species with larger habitat breadth do have more chances to escape from multiple pressure types and are consistently less threatened across taxa and spatial settings.

Although almost half of all measurements of body size were significant, the meta-analyses revealed that the relationship between body size and extinction risk is not unidirectional. The interplay between body size and threat type is one of the reasons for this phenomenon. While larger bird species are threatened by overexploitation, smaller bird species are threatened by habitat loss or degradation (Owens and Bennet, 2000). The same trend seems to apply at least to marine fishes (Olden, Hogan & Zanden, 2007) and mammals (González-Suárez *et al.*, 2013), taxa that are often targeted directly and selectively by man. Independently of threats, relationships may not even be linear. Threatened freshwater fishes are found both at the smaller and larger spectrum of body sizes (Olden *et al.*, 2007), and the same bimodal relationship is found when pooling all vertebrates together (Ripple *et al.*, 2017). In general, this bimodality seems to be derived from threat type, with different threats leading to increasing extinction risk of different body size classes.

Other traits for which we could not perform a quantitative analysis have also shown to be useful in predicting extinction risk under certain circumstances, such as those traits related to speed of life cycle and reproductive output. Threat status has been positively related to species with decreased fecundity (Cardillo, 2003; González-Suárez & Revilla, 2013; Böhm *et al.*, 2016; Ribeiro *et al.*, 2016; but see Pinsky & Byler, 2015; Sreekar *et al.*, 2015), larger egg/neonatal sizes (Cardillo *et al.*, 2005; Jones, Fielding & Sullivan, 2006; González-Suárez & Revilla, 2013; Pinsky & Byler, 2015) and longer generation lengths (Anderson *et al.*, 2011; Hanna & Cardillo, 2013; Jeppsson & Forslund, 2014; Comeros-Raynal *et al.*, 2016; but see Chessman, 2013. These traits usually correlate with each other and with body size and longevity: bigger, longer-lived species often have lower fecundity, bigger egg/neonatal sizes and longer generation lengths. These traits reduce the capability of species to compensate for high mortality rates (Pimm, Jones & Diamond, 1988; Purvis *et al.*, 2000a; González-Suárez *et al.*, 2013), even if their longer longevities should make them more apt to resist at lower densities as they survive longer and might be able to overcome short-lived threats (Pimm *et al.*, 1988). When species are directly persecuted by man, they are often bigger, with larger fecundity and egg/neonatal sizes (Owens & Bennett, 2000; González-Suárez *et al.*, 2013), and longer longevity alone is not sufficient to compensate for the high mortality. But when the threat is habitat loss, which indirectly increases mortality and/or reduces natality rates, the trend is non-existent or even reversed (Owens & Bennett, 2000; González-Suárez *et al.*, 2013), this being possibly due to the advantages of longer longevity alone.

Traits indicating preference towards specific environmental niches are commonly used across taxa and many data are available about them. Among those, temperature (optimal temperature or temperature of the species across its geographical range) and temperature range (range of temperatures tolerated by the species or range of temperatures found across its geographical range) were often important predictors in the studies that used them. Species with lower average temperatures within their range or narrower temperature ranges are especially at risk due to an increasingly warmer climate (Jiguet *et al.*, 2010; Grenouillet & Comte, 2014; Flousek *et al.*, 2015). In contrast, thriving under broad temperature ranges grants species the necessary flexibility to deal with environmental or climatic change and hence lower their extinction risk (Chessman, 2013; Lootvoet, Philippon & Bessa-Gomes, 2015). When exceptions were found, these were due to the correlation of temperature with the true causes of change in extinction risk (e.g. Cooper *et al.*, 2008).

Although the generality of the pattern could not be confirmed across studies, species depending on habitats more affected by human influence are often more threatened (Stefanaki *et al.*, 2015; Powney *et al.*, 2014). In Greece, flowering plants occurring in coastal or ruderal habitats, under pressure from urbanization and tourism, were more at risk than flowering plants occurring on cliffs or high-mountain vegetation, the latter habitats being under lower human pressure (Stefanaki *et al.*, 2015). British plant species with lower affinity to nitrogen-rich soils are declining due to the intensification of agriculture, which has led to increased inputs of nitrogen in otherwise nitrogen-poor soils (Powney *et al.*, 2014). Likewise, microhabitat type was a good predictor of extinction risk in some studies due to some microhabitats becoming rarer with increased human pressure (Parent & Schriml, 1995; Seibold *et al.*, 2015). A striking example is the decline of saproxylic beetles that use dead wood of large diameter in Germany, as forest management options often lead to the scarcity of such microhabitat (Seibold *et al.*, 2015). These observations give support to recent claims that predicting extinction risk requires considering the threat type and using different variables related to human use of species and habitats (Murray *et al.*, 2014).

Both diet breadth and type were significant predictors across several studies. The diet of a species can be important in leading to and predicting extinction in two ways. Species restricted to fewer dietary options have shown to be more threatened (Basset *et al.*, 2015; Jeppsson & Forslund, 2014; González-Suárez *et al.*, 2013; Matsuzaki *et al.*, 2011; Mattila *et al.*, 2008), probably due to lower flexibility in switching to other options when the availability of their preferred food source decreases (Purvis, Jones, & Mace, 2000). On the other hand, diet type, namely the trophic position of a species, may be as important. Species at higher trophic levels tend to be more threatened (Chessman, 2013; Bender *et al.*, 2013; Cardillo *et al.*, 2004; Purvis *et al.*, 2000a) and often provide early warnings of extinction across the entire food chain (Cardoso *et al.*, 2010). The greater dependence on the densities and larger foraging areas of prey species may lead to such a pattern (Carbone & Gittleman, 2002), with synergistic effects between resource abundance and other factors contributing to the decline of, for example, predators. With the density of wildlife dwindling everywhere (e.g. Hallmann *et al.*, 2017), and everything else being equal, top predators are expected to be more at risk.

Migration distance was often tested and found to be an important predictor. Most studies on migration distance are of birds. Long distance migrants tend to be more at risk, which could be either due to phenological mismatch due to climate change (Amano & Yamaura, 2007; Jiguet *et al.*, 2010; Thaxter *et al.*, 2010; Flousek *et al.*, 2015), dependence on the good quality of at least two habitats or sites (Jiguet *et al.*, 2010; Flousek *et al.*, 2015), or to increased competition with resident species that, in temperate regions, survive through increasingly less severe winters (Jiguet *et al.*, 2010; Amano & Yamaura, 2007).

Finally, there are also traits that were found to be significant but only studied for one or two taxa. These include a wide array of morphological traits that are taxon-specific. Some plant growth forms (e.g. herbaceous, bush or tree) are more threatened than others. Perennial growth forms can sustain populations through harsh times (Stefanaki *et al.*, 2015) but might be more affected by forest loss (Leão *et al.*, 2014). Mammals going through a hibernating or torpor phase are less prone to becoming extinct, due to a greater capacity to avoid harsher seasonal conditions (Liow *et al.*, 2009). The life stage in which an insect overwinters (egg, larva, pupa or adult) influences vulnerability (e.g. Powney *et al.*, 2015; Jeppsson & Forslund, 2014; Mattila *et al.*, 2009). At least for some studies with applied relevance, Cardillo & Meijaard (2012) claim that ‘researchers should adopt a somewhat “smaller picture” view by restricting the geographical and taxonomic scope of comparative analyses, and aiming for clearer, more focused outcomes on particular hypotheses’. We corroborate that restricting the studies in these two dimensions might prove useful when the goal goes beyond understanding the general pattern and requires true predictive power for species extinctions.

### (2) Generalization

Given the inherent bias of past studies, any generalizations require critical consideration. Geographical range and habitat breadth seem to be very well supported across taxa and regions, even if most past studies using such traits were on vertebrates. Both are consistently negatively related to extinction risk and might be seen as representing a single phenomenon: the range or rarity of a species in two different dimensions (area and habitat). Species with larger ranges, be these spatial or biotic, have more chance of surviving in case of diminishing availability of resources, and the risk of their populations or the entire species vanishing is smaller. These traits can therefore be confidently used as predictors of extinction risk across taxa. Area and habitat are in fact two of the three dimensions of rarity preconized by Rabinowitz (1981): geographical range size, habitat breadth, and local abundance. The latter was seldom used probably due to the scarcity of abundance data for most taxa (the Prestonian shortfall, Cardoso *et al.*, 2011b) but is certainly crucial to fully understand the extinction phenomenon.

Body size, on the other hand, seems to be at least taxon dependent, probably because, as previously mentioned, it represents different ways in which species interact with their environment and therefore how they affect their risk of extinction (González-Suárez *et al.*, 2013; Ripple *et al.*, 2017). This trait is often studied as a proxy for traits that may be very hard to measure or are very abstract. If for animals it usually is related to resource availability, as larger animals require more, often scarce, resources, being these, space, food or other, for plants it represents competitive ability, with larger plants being able to better exploit, for example, the sun, by growing taller and overshadowing smaller species, or water and mineral resources found deeper underground.

In this review, we reinforced the notion that species with smaller ranges, and those with narrow habitat breadths are more at risk than others, regardless of the taxon or geographic distribution. We must emphasize, however, that we still lack a complete and unbiased picture of the relation between traits and extinction risk and that future studies could and should provide insights much beyond what is possible now. Many traits were found to be important across studies but have seldom been studied or are relevant for only some taxa. Not only that, but the intricate links between e.g. body size and extinctions provide reason for further studies to focus not only on the threat status of a species, but also on the underlying threat (whether it be human persecution, habitat degradation, climate change, or invasive species).

## VI. Acknowledgements

We thank Sini Seppälä, Jon Rikberg and Fernando Urbano-Tenorio for their suggestions in earlier stages of this work, and to Marina Ferreira for useful suggestions on the manuscript. We would also like to thank Cathryn Primrose-Mathisen who provided professional English language assistance during the preparation of this article. She was not responsible for reviewing the final version. F.C. and this project were funded by Kone Foundation.

## Reference

Amano, T., & Yamaura, Y. (2007). Ecological and life-history traits related to range contractions among breeding birds in Japan. Biological Conservation, 137(2), 271–282. doi:10.1016/j.biocon.2007.02.010

Anderson, S. C., Farmer, R. G., Ferretti, F., Houde, A. L. S., & Hutchings, J. A. (2011). Correlates of Vertebrate Extinction Risk in Canada. BioScience, 61(7), 538–549. doi:10.1525/bio.2011.61.7.8

Arbetman, M. P., Gleiser, G., Morales, C. L., Williams, P., & Aizen, M. A. (2017). Global decline of bumblebees is phylogenetically structured and inversely related to species range size and pathogen incidence. Proceedings of the Royal Society B: Biological Sciences, 284(1859), 20170204. doi:10.1098/rspb.2017.0204

Basset, Y., Barrios, H., Segar, S., Srygley, R. B., Aiello, A., Warren, A. D., … Ramirez, J. A. (2015). The Butterflies of Barro Colorado Island, Panama: Local Extinction since the 1930s. PLoS ONE, 10(8), e0136623. doi:10.1371/journal.pone.0136623

Bender, M. g., Floeter, S. r., Mayer, F. p., Vila-Nova, D. a., Longo, G. o., Hanazaki, N., … Ferreira, C. e. l. (2013). Biological attributes and major threats as predictors of the vulnerability of species: a case study with Brazilian reef fishes. Oryx, 47(02), 259–265. doi:10.1017/S003060531100144X

Bland, L. M., Collen, B., Orme, C. D. L., & Bielby, J. (2015). Predicting the conservation status of data-deficient species. Conservation Biology, 29(1), 250–259. doi:10.1111/cobi.12372

Böhm, M., Williams, R., Bramhall, H. R., McMillan, K. M., Davidson, A. D., Garcia, A., … Collen, B. (2016). Correlates of extinction risk in squamate reptiles: the relative importance of biology, geography, threat and range size. Global Ecology and Biogeography, 25(4), 391–405. doi:10.1111/geb.12419

Burgman, M. A. (1989). The Habitat Volumes of Scarce and Ubiquitous Plants: A Test of the Model of Environmental Control. The American Naturalist, 133(2), 228–239. doi:10.1086/284912

Carbone, C., & Gittleman, J. L. (2002). A Common Rule for the Scaling of Carnivore Density. Science, 295(5563), 2273–2276. doi:10.1126/science.1067994

Cardillo, M. (2003). Biological determinants of extinction risk: why are smaller species less vulnerable? Animal Conservation, 6(1), 63–69. doi:10.1017/S1367943003003093

Cardillo, M., Mace, G. M., Gittleman, J. L., Jones, K. E., Bielby, J., & Purvis, A. (2008). The predictability of extinction: biological and external correlates of decline in mammals. Proceedings of the Royal Society of London B: Biological Sciences, 275(1641), 1441–1448. doi:10.1098/rspb.2008.0179

Cardillo, M., Mace, G. M., Jones, K. E., Bielby, J., Bininda-Emonds, O. R. P., Sechrest, W., … Purvis, A. (2005). Multiple Causes of High Extinction Risk in Large Mammal Species. Science, 309(5738), 1239–1241. doi:10.1126/science.1116030

Cardillo, M., & Meijaard, E. (2012). Are comparative studies of extinction risk useful for conservation? Trends in Ecology & Evolution, 27(3), 167–171. doi:10.1016/j.tree.2011.09.013

Cardillo, M., Purvis, A., Sechrest, W., Gittleman, J. L., Bielby, J., & Mace, G. M. (2004). Human Population Density and Extinction Risk in the World’s Carnivores. PLoS Biol, 2(7), e197. doi:10.1371/journal.pbio.0020197

Cardoso, P., Borges, P. A. V., Triantis, K. A., Ferrández, M. A., & Martín, J. L. (2011). Adapting the IUCN Red List criteria for invertebrates. Biological Conservation, 144(10), 2432–2440. doi:10.1016/j.biocon.2011.06.020

Cardoso, P., Borges, P. A. V., Triantis, K. A., Ferrández, M. A., & Martín, J. L. (2012). The underrepresentation and misrepresentation of invertebrates in the IUCN Red List. Biological Conservation, 149(1), 147–148. doi:10.1016/j.biocon.2012.02.011

Cardoso, P., Erwin, T. L., Borges, P. A. V., & New, T. R. (2011). The seven impediments in invertebrate conservation and how to overcome them. Biological Conservation, 144(11), 2647–2655. doi:10.1016/j.biocon.2011.07.024

Chaplin-Kramer, R., O’Rourke, M. E., Blitzer, E. J., & Kremen, C. (2011). A meta-analysis of crop pest and natural enemy response to landscape complexity. Ecology Letters, 14(9), 922–932. doi:10.1111/j.1461-0248.2011.01642.x

Chessman, B. C. (2013). Identifying species at risk from climate change: Traits predict the drought vulnerability of freshwater fishes. Biological Conservation, 160, 40–49. doi:10.1016/j.biocon.2012.12.032

Clavel, J., Julliard, R., & Devictor, V. (2011). Worldwide decline of specialist species: toward a global functional homogenization? Frontiers in Ecology and the Environment, 9(4), 222–228. doi:10.1890/080216

Comeros-Raynal, M. T., Polidoro, B. A., Broatch, J., Mann, B. Q., Gorman, C., Buxton, C. D., … Carpenter, K. E. (2016). Key predictors of extinction risk in sea breams and porgies (Family: Sparidae). Biological Conservation, 202, 88–98. doi:10.1016/j.biocon.2016.08.027

Cooper, N., Bielby, J., Thomas, G. H., & Purvis, A. (2008). Macroecology and extinction risk correlates of frogs. Global Ecology and Biogeography, 17(2), 211–221. doi:10.1111/j.1466-8238.2007.00355.x

DeAngelis, D. L., & Grimm, V. (2014). Individual-based models in ecology after four decades. F1000Prime Reports, 6. doi:10.12703/P6-39

Fisher, D. O., & Owens, I. P. F. (2004). The comparative method in conservation biology. Trends in Ecology & Evolution, 19(7), 391–398. doi:10.1016/j.tree.2004.05.004

Flousek, J., Telenský, T., Hanzelka, J., & Reif, J. (2015). Population Trends of Central European Montane Birds Provide Evidence for Adverse Impacts of Climate Change on High-Altitude Species. PLoS ONE, 10(10), e0139465. doi:10.1371/journal.pone.0139465

Gaston, K. J., Blackburn, T. M., Greenwood, J. J. D., Gregory, R. D., Quinn, R. M., & Lawton, J. H. (2002). Abundance–occupancy relationships. Journal of Applied Ecology, 37(s1), 39–59. doi:10.1046/j.1365-2664.2000.00485.x

González-Suárez, M., Gómez, A., & Revilla, E. (2013). Which intrinsic traits predict vulnerability to extinction depends on the actual threatening processes. Ecosphere, 4(6), 1–16. doi:10.1890/ES12-00380.1

González-Suárez, M., & Revilla, E. (2013). Variability in life-history and ecological traits is a buffer against extinction in mammals. Ecology Letters, 16(2), 242–251. doi:10.1111/ele.12035

Grenouillet, G., & Comte, L. (2014). Illuminating geographical patterns in species’ range shifts. Global Change Biology, 20(10), 3080–3091. doi:10.1111/gcb.12570

Hallmann, C. A., Sorg, M., Jongejans, E., Siepel, H., Hofland, N., Schwan, H., … Kroon, H. de. (2017). More than 75 percent decline over 27 years in total flying insect biomass in protected areas. PLoS ONE, 12(10), e0185809. doi:10.1371/journal.pone.0185809

Hanna, E., & Cardillo, M. (2013). A comparison of current and reconstructed historic geographic range sizes as predictors of extinction risk in Australian mammals. Biological Conservation, 158, 196–204. doi:10.1016/j.biocon.2012.08.014

IUCN. (2019). The IUCN Red List of Threatened Species. Version 2018-2. Retrieved January 16, 2019, from http://www.iucnredlist.org

Jeppsson, T., & Forslund, P. (2014). Species’ traits explain differences in Red list status and long-term population trends in longhorn beetles: Traits and extinction risk in longhorn beetles. Animal Conservation, 17(4), 332–341. doi:10.1111/acv.12099

Jiguet, F., Gregory, R. D., Devictor, V., Green, R. E., Voříšek, P., Van Strien, A., & Couvet, D. (2010). Population trends of European common birds are predicted by characteristics of their climatic niche. Global Change Biology, 16(2), 497–505. doi:10.1111/j.1365-2486.2009.01963.x

Jones, M. J., Fielding, A., & Sullivan, M. (2006). Analysing Extinction Risk in Parrots using Decision Trees. Biodiversity & Conservation, 15(6), 1993–2007. doi:10.1007/s10531-005-4316-1

Koh, L. P., Sodhi, N. S., & Brook, B. W. (2004). Ecological Correlates of Extinction Proneness in Tropical Butterflies: Extinction Correlates of Tropical Butterflies. Conservation Biology, 18(6), 1571–1578. doi:10.1111/j.1523-1739.2004.00468.x

Leão, T. C. C., Fonseca, C. R., Peres, C. A., & Tabarelli, M. (2014). Predicting Extinction Risk of Brazilian Atlantic Forest Angiosperms: Neotropical Plant Extinction Risk. Conservation Biology, 28(5), 1349–1359. doi:10.1111/cobi.12286

Liow, L. H., Fortelius, M., Lintulaakso, K., Mannila, H., & Stenseth, N. C. (2009). Lower Extinction Risk in Sleep‐or‐Hide Mammals. The American Naturalist, 173(2), 264–272. doi:10.1086/595756

Lootvoet, A. C., Philippon, J., & Bessa-Gomes, C. (2015). Behavioral Correlates of Primates Conservation Status: Intrinsic Vulnerability to Anthropogenic Threats. PLoS ONE, 10(10), e0135585. doi:10.1371/journal.pone.0135585

Luiz, O. J., Woods, R. M., Madin, E. M. P., & Madin, J. S. (2016). Predicting IUCN Extinction Risk Categories for the World’s Data Deficient Groupers (Teleostei: Epinephelidae). Conservation Letters, 9(5), 342–350. doi:10.1111/conl.12230

Matsuzaki, S.-I. S., Takamura, N., Arayama, K., Tominaga, A., Iwasaki, J., & Washitani, I. (2011). Potential impacts of non-native channel catfish on commercially important species in a Japanese lake, as inferred from long-term monitoring data. Aquatic Conservation: Marine and Freshwater Ecosystems, 21(4), 348–357. doi:10.1002/aqc.1198

Mattila, N., Kotiaho, J. S., Kaitala, V., & Komonen, A. (2008). The use of ecological traits in extinction risk assessments: A case study on geometrid moths. Biological Conservation, 141(9), 2322–2328. doi:10.1016/j.biocon.2008.06.024

McKinney, M. L. (1997). Extinction Vulnerability and Selectivity: Combining Ecological and Paleontological Views. Annual Review of Ecology and Systematics, 28, 495–516. Retrieved from http://www.jstor.org/stable/2952502

McKinney, M. L., & Lockwood, J. L. (1999). Biotic homogenization: a few winners replacing many losers in the next mass extinction. Trends in Ecology & Evolution, 14(11), 450–453. doi:10.1016/S0169-5347(99)01679-1

Murray, K. A., Verde Arregoitia, L. D., Davidson, A., Di Marco, M., & Di Fonzo, M. M. I. (2014). Threat to the point: improving the value of comparative extinction risk analysis for conservation action. Global Change Biology, 20(2), 483–494. doi:10.1111/gcb.12366

Olden, J. D., Hogan, Z. S., & Zanden, M. J. V. (2007). Small fish, big fish, red fish, blue fish: size-biased extinction risk of the world’s freshwater and marine fishes. Global Ecology and Biogeography, 16(6), 694–701. doi:10.1111/j.1466-8238.2007.00337.x

Olson, D. M., Dinerstein, E., Wikramanayake, E. D., Burgess, N. D., Powell, G. V. N., Underwood, E. C., … Kassem, K. R. (2001). Terrestrial Ecoregions of the World: A New Map of Life on Earth: A new global map of terrestrial ecoregions provides an innovative tool for conserving biodiversity. BioScience, 51(11), 933–938. doi:10.1641/0006-3568(2001)051[0933:TEOTWA]2.0.CO;2

Owens, I. P. F., & Bennett, P. M. (2000). Ecological basis of extinction risk in birds: Habitat loss versus human persecution and introduced predators. Proceedings of the National Academy of Sciences, 97(22), 12144–12148. doi:10.1073/pnas.200223397

Parent, S., & Schriml, L. M. (1995). A model for the determination of fish species at risk based upon life-history traits and ecological data. Canadian Journal of Fisheries and Aquatic Sciences, 52(8), 1768–1781. doi:10.1139/f95-769

Pimm, S. L., Jones, H. L., & Diamond, J. (1988). On the Risk of Extinction. The American Naturalist, 132(6), 757–785. Retrieved from http://www.jstor.org/stable/2462261

Pinsky, M. L., & Byler, D. (2015). Fishing, fast growth and climate variability increase the risk of collapse. Proceedings of the Royal Society B: Biological Sciences, 282(1813), 20151053. doi:10.1098/rspb.2015.1053

Polaina, E., Revilla, E., & González-Suárez, M. (2016). Putting susceptibility on the map to improve conservation planning, an example with terrestrial mammals. Diversity and Distributions, 22(8), 881–892. doi:10.1111/ddi.12452

Powney, G. D., Rapacciuolo, G., Preston, C. D., Purvis, A., & Roy, D. B. (2014). A phylogenetically-informed trait-based analysis of range change in the vascular plant flora of Britain. Biodiversity and Conservation, 23(1), 171–185. doi:10.1007/s10531-013-0590-5

Prugh, L. R. (2009). An evaluation of patch connectivity measures. Ecological Applications, 19(5), 1300–1310. doi:10.1890/08-1524.1

Purvis, A., Gittleman, J. L., Cowlishaw, G., & Mace, G. M. (2000). Predicting extinction risk in declining species. Proceedings of the Royal Society of London B: Biological Sciences, 267(1456), 1947–1952. doi:10.1098/rspb.2000.1234

Purvis, A., Jones, K. E., & Mace, G. M. (2000). Extinction. BioEssays, 22(12), 1123–1133. doi:10.1002/1521-1878(200012)22:12<1123::AID-BIES10>3.0.CO;2-C

Rabinowitz, D. (1981). Seven forms of Rarity. In Biological aspects of rare plant conservation (pp. 205–217). New York: John Wiley & Sons Ltd.

Ribeiro, J., Colli, G. R., Caldwell, J. P., & Soares, A. M. V. M. (2016). An integrated trait-based framework to predict extinction risk and guide conservation planning in biodiversity hotspots. Biological Conservation, 195, 214–223. doi:10.1016/j.biocon.2015.12.042

Ripple, W. J., Wolf, C., Newsome, T. M., Hoffmann, M., Wirsing, A. J., & McCauley, D. J. (2017). Extinction risk is most acute for the world’s largest and smallest vertebrates. Proceedings of the National Academy of Sciences, 201702078. doi:10.1073/pnas.1702078114

Rosenthal, R. (1991). Meta-Analytic Procedures for Social Research (Vol. 6). London: SAGE Publications.

Seibold, S., Brandl, R., Buse, J., Hothorn, T., Schmidl, J., Thorn, S., & Müller, J. (2015). Association of extinction risk of saproxylic beetles with ecological degradation of forests in Europe: Beetle Extinction and Forest Degradation. Conservation Biology, 29(2), 382–390. doi:10.1111/cobi.12427

Slatyer, R. A., Hirst, M., & Sexton, J. P. (2013). Niche breadth predicts geographical range size: a general ecological pattern. Ecology Letters, 16(8), 1104–1114. doi:10.1111/ele.12140

Sodhi, N. S., Koh, L. P., Peh, K. S.-H., Tan, H. T. W., Chazdon, R. L., Corlett, R. T., … Bradshaw, C. J. A. (2008). Correlates of extinction proneness in tropical angiosperms. Diversity and Distributions, 14(1), 1–10. doi:10.1111/j.1472-4642.2007.00398.x

Sreekar, R., Huang, G., Zhao, J.-B., Pasion, B. O., Yasuda, M., Zhang, K., … Harrison, R. D. (2015). The use of species-area relationships to partition the effects of hunting and deforestation on bird extirpations in a fragmented landscape. Diversity and Distributions, 21(4), 441–450. doi:10.1111/ddi.12292

Stefanaki, A., Kantsa, A., Tscheulin, T., Charitonidou, M., & Petanidou, T. (2015). Lessons from Red Data Books: Plant Vulnerability Increases with Floral Complexity. PLoS ONE, 10(9), e0138414. doi:10.1371/journal.pone.0138414

Stuart, S. N., Wilson, E. O., McNeely, J. A., Mittermeier, R. A., & Rodríguez, J. P. (2010). The Barometer of Life. Science, 328(5975), 177–177. doi:10.1126/science.1188606

Sullivan, M. S., Gilbert, F., Rotheray, G., Croasdale, S., & Jones, M. (2000). Comparative analyses of correlates of Red data book status: a case study using European hoverflies (Diptera: Syrphidae). Animal Conservation, 3(2), 91–95.

Terzopoulou, S., Rigal, F., Whittaker, R. J., Borges, P. A. V., & Triantis, K. A. (2015). Drivers of extinction: the case of Azorean beetles. Biology Letters, 11(6), 20150273. doi:10.1098/rsbl.2015.0273

Thaxter, C. B., Joys, A. C., Gregory, R. D., Baillie, S. R., & Noble, D. G. (2010). Hypotheses to explain patterns of population change among breeding bird species in England. Biological Conservation, 143(9), 2006–2019. doi:10.1016/j.biocon.2010.05.004

Verde Arregoitia, L. D. (2016). Biases, gaps, and opportunities in mammalian extinction risk research. Mammal Review, 46(1), 17–29. doi:10.1111/mam.12049

Viechtbauer, W. (2010). Conducting Meta-Analyses in R with the metafor Package. Journal of Statistical Software, 36(3), 1–48. doi:10.18637/jss.v036.i03

Wang, Y., Si, X., Bennett, P. M., Chen, C., Zeng, D., Zhao, Y., … Ding, P. (2018). Ecological correlates of extinction risk in Chinese birds. Ecography, 41(5), 782–794. doi:10.1111/ecog.03158

