## Appendix S1 & S3-4 for "A review of the relation between species traits and extinction risk"

### VIII. Supplementary materials

Appendix S1 – List of all studies and traits included in the analyses

Table S1: List of all traits (after joining together traits with different names but depicting a similar phenomenon), their category, as well as the number of studies and measurements (the number of measurements corresponds to the total number of statistical coefficients of each trait, usually corresponding to the number of statistical tests for that trait) for each of the traits, and the reference in which they appear (see below). **CSR:** C – S – R Triangle theory (Competitive – Stress tolerant – Ruderal).

| Trait | Number of studies | Number of measurements | Reference Number |
|---|---|---|---|
| Accessibility | 2 | 25 | 3, 16 |
| Active dispersion | 1 | 8 | 120 |
| Age at dispersal | 1 | 2 | 69 |
| Age at eye opening | 3 | 21 | 25, 26, 69 |
| Air-breathing capability | 1 | 2 | 59 |
| Altitude | 16 | 102 | 3, 15, 16, 20, 54, 66, 85, 90, 98, 100, 104, 107, 150, 152, 159, 169 |
| Altitude range | 7 | 37 | 54, 85, 98, 100, 106, 123, 155 |
| Altitudinal range shift | 1 | 2 | 50 |
| Ambush predation | 1 | 2 | 141 |
| Amount of floral reward | 1 | 4 | 62 |
| Annual productivity range | 1 | 1 | 38 |
| Body armour | 1 | 6 | 20 |
| Body shape | 2 | 10 | 66, 171 |
| Body size | 141 | 700 | 1, 2, 3, 4, 7, 8, 9, 11, 12, 15, 16, 20, 21, 23, 24, 25, 26, 27, 28, 29, 30, 31, 32, 33, 34, 35, 38, 39, 40, 42, 43, 44, 45, 48, 49, 51, 53, 54, 55, 56, 57, 58, 59, 60, 61, 63, 64, 65, 66, 67, 68, 69, 70, 71, 72, 74, 75, 76, 77, 78, 79, 80, 81, 82, 83, 84, 85, 86, 87, 88, 89, 90, 91, 92, 93, 94, 95, 96, 98, 99, 100, 101, 102, 103, 104, 105, 106, 107, 108, 109, 110, 111, 112, 113, 114, 116, 117, 119, 120, 121, 122, 123, 124, 125, 126, 127, 128, 129, 130, 131, 132, 133, 134, 135, 136, 137, 138, 139, 140, 141, 142, 143, 144, 146, 147, 148, 151, 152, 154, 156, 157, 158, 161, 162, 163, 164, 168, 169, 171, 172, 173 |
| Body size at weaning | 2 | 3 | 39, 69 |
| Body size variability | 1 | 22 | 64 |
| Brain size | 6 | 53 | 1, 65, 134, 154, 162, 168 |
| Change in human population density | 3 | 21 | 25, 26, 102 |
| Chela shape | 1 | 18 | 15 |
| Chela size | 1 | 18 | 15 |
| Chemical defence | 1 | 3 | 6 |
| Circadian period | 17 | 63 | 25, 26, 27, 38, 40, 42, 69, 70, 96, 98, 100, 106, 120, 130, 140, 143, 164 |
| Circadian range | 1 | 4 | 93 |
| Climate range | 2 | 8 | 18, 169 |
| Clutch volume | 1 | 2 | 90 |
| Colour detectability category | 2 | 4 | 10, 86 |
| Colour pattern variability | 1 | 4 | 14 |
| Competitor richness | 4 | 30 | 59, 82, 86, 120 |
| Compound metrics | 6 | 66 | 9, 46, 87, 98, 148, 169 |

| | | | |
|---|---|---|---|
| Considered a pest | 1 | 10 | 120 |
| Controlled by man | 1 | 10 | 120 |
| Corolla segmentation | 1 | 3 | 159 |
| Country responsible for management | 1 | 4 | 131 |
| CSR competitiveness | 1 | 3 | 167 |
| CSR ruderality | 2 | 6 | 52, 167 |
| CSR strategy | 1 | 2 | 91 |
| CSR stress tolerance | 1 | 3 | 167 |
| Depending on symbioses | 1 | 23 | 120 |
| Development type | 13 | 57 | 3, 7, 16, 55, 70, 72, 88, 98, 119, 120, 141, 163, 168 |
| Diaspore mass | 1 | 3 | 92 |
| Diaspore type | 2 | 6 | 62, 91 |
| Diet breadth | 35 | 153 | 4, 8, 9, 10, 14, 19, 25, 26, 39, 43, 52, 56, 63, 69, 71, 76, 86, 88, 93, 96, 98, 106, 108, 110, 111, 112, 120, 123, 125, 130, 139, 141, 148, 158, 170 |
| Diet type | 49 | 236 | 4, 8, 9, 10, 11, 16, 25, 26, 27, 29, 36, 38, 39, 40, 47, 49, 59, 67, 69, 77, 78, 79, 82, 84, 90, 95, 98, 99, 102, 103, 106, 112, 120, 121, 125, 127, 129, 130, 131, 132, 140, 141, 142, 152, 157, 158, 163, 168, 173 |
| Direct exploitation | 13 | 65 | 3, 11, 21, 34, 48, 77, 83, 102, 106, 120, 131, 132, 173 |
| Dispersal ability | 21 | 95 | 9, 13, 20, 32, 45, 46, 56, 66, 77, 110, 118, 120, 125, 128, 129, 147, 162, 167, 168, 172, 173 |
| Dispersal agent | 8 | 36 | 94, 120, 147, 150, 156, 165, 166, 171 |
| Duration of flight period | 8 | 22 | 9, 14, 22, 76, 110, 111, 137, 160 |
| Duration of reproductive season | 10 | 34 | 8, 29, 66, 71, 83, 88, 91, 127, 159, 168 |
| Economic value | 1 | 2 | 61 |
| Egg adhesion | 1 | 4 | 29 |
| Egg buoyancy | 1 | 4 | 29 |
| Egg/neonatal size | 21 | 81 | 3, 25, 26, 43, 45, 46, 64, 66, 69, 82, 87, 90, 101, 117, 119, 127, 131, 132, 134, 141, 168 |
| Egg/neonatal size variability | 1 | 8 | 64 |
| Egg-laying behaviour | 1 | 3 | 86 |
| Endemism | 4 | 25 | 44, 45, 94, 120 |
| Endotherm | 1 | 10 | 120 |
| Evapotranspiration | 6 | 41 | 25, 26, 33, 35, 98, 169 |
| Evolutionary plasticity | 2 | 18 | 82, 120 |
| Fecundity | 73 | 434 | 2, 3, 7, 8, 9, 12, 13, 15, 16, 23, 25, 26, 27, 29, 32, 33, 35, 39, 43, 46, 49, 51, 52, 55, 56, 63, 64, 65, 66, 69, 70, 71, 74, 75, 77, 78, 81, 82, 83, 87, 88, 90, 95, 96, 98, 101, 102, 106, 117, 119, 120, 123, 125, 127, 128, 130, 131, 132, 133, 134, 140, 141, 142, 143, 147, 154, 155, 158, 160, 162, 163, 168, 173 |
| Fecundity variability | 2 | 22 | 64, 168 |
| Fertilization mode | 8 | 43 | 62, 90, 118, 150, 156, 166, 171, 172 |
| Fledgling duration | 4 | 13 | 71, 90, 128, 134 |

| | | | |
|---|---|---|---|
| Flight period to adult life span | 1 | 2 | 22 |
| Floral colour | 2 | 7 | 150, 159 |
| Floral display | 1 | 3 | 20 |
| Floral shape | 1 | 10 | 159 |
| Floral size | 1 | 6 | 159 |
| Forearm length | 1 | 8 | 64 |
| Forearm length variability | 1 | 8 | 64 |
| Fruit dehiscence | 2 | 7 | 20, 62 |
| Functional reproductive unit | 1 | 4 | 159 |
| Generation length | 41 | 239 | 3, 4, 25, 26, 27, 29, 32, 33, 34, 55, 61, 63, 64, 66, 69, 71, 74, 75, 76, 77, 78, 81, 83, 88, 93, 95, 96, 101, 107, 108, 120, 125, 126, 127, 128, 130, 132, 139, 140, 142, 168 |
| Generation length variability | 1 | 17 | 64 |
| Geographic origin | 2 | 19 | 91, 120 |
| Geographical range size | 81 | 377 | 2, 4, 5, 9, 10, 11, 14, 15, 16, 18, 20, 25, 26, 27, 30, 32, 33, 34, 35, 37, 38, 39, 40, 43, 44, 48, 52, 53, 54, 56, 59, 64, 66, 67, 69, 70, 72, 78, 81, 82, 85, 86, 87, 89, 95, 97, 98, 99, 100, 102, 104, 105, 106, 107, 117, 119, 120, 123, 130, 133, 135, 137, 139, 140, 142, 143, 144, 149, 150, 152, 153, 155, 156, 157, 159, 161, 164, 168, 171, 172, 173 |
| Geographical range size change | 3 | 4 | 56, 72, 123 |
| Geographical range size of host | 1 | 1 | 111 |
| Gestation/incubation length | 19 | 117 | 3, 25, 26, 27, 53, 63, 64, 65, 66, 69, 71, 81, 87, 90, 96, 103, 134, 139, 140 |
| Gestation/incubation length variability | 1 | 8 | 64 |
| Gonad size | 5 | 20 | 102, 114, 115, 117, 119 |
| Gross domestic product | 2 | 10 | 139, 163 |
| Habitat breadth | 54 | 258 | 2, 4, 15, 16, 18, 25, 26, 33, 35, 43, 46, 49, 54, 56, 61, 63, 69, 71, 77, 84, 85, 87, 89, 90, 93, 94, 95, 98, 102, 104, 106, 110, 122, 123, 126, 130, 133, 137, 138, 139, 141, 142, 146, 149, 150, 153, 154, 158, 160, 161, 164, 168, 172, 173 |
| Habitat type | 56 | 364 | 2, 4, 7, 14, 15, 17, 20, 21, 34, 38, 39, 40, 42, 44, 45, 48, 52, 54, 55, 59, 69, 70, 72, 77, 78, 79, 85, 86, 88, 89, 91, 93, 95, 97, 100, 105, 106, 119, 120, 127, 129, 130, 134, 135, 142, 143, 150, 151, 153, 156, 157, 158, 159, 162, 164, 168 |
| Historical geographical range size | 4 | 10 | 45, 49, 69, 163 |
| Home range size | 17 | 81 | 13, 23, 25, 26, 27, 33, 38, 40, 64, 67, 69, 84, 96, 103, 130, 139, 140 |
| Home range size variability | 1 | 8 | 64 |
| Human footprint | 7 | 34 | 16, 37, 39, 43, 98, 103, 164 |
| Human night-time lights | 1 | 1 | 38 |
| Human population density | 14 | 167 | 15, 16, 25, 26, 27, 33, 37, 38, 84, 85, 139, 163, 164, 169 |

| | | | |
|---|---|---|---|
| Human resource intake | 2 | 90 | 15, 85 |
| Humidity | 1 | 1 | 72 |
| Incubating sex | 1 | 1 | 128 |
| Insularity | 13 | 83 | 16, 25, 26, 27, 37, 38, 40, 81, 82, 140, 155, 163, 167 |
| Interbirth interval | 11 | 63 | 25, 26, 27, 33, 55, 64, 69, 73, 130, 139, 140 |
| Interbirth interval variability | 1 | 8 | 64 |
| Internal or external fertilization | 1 | 1 | 72 |
| Investment in reproduction | 3 | 11 | 4, 73, 90 |
| Land-use change | 4 | 16 | 37, 43, 102, 164 |
| Land-use proportion | 11 | 102 | 15, 16, 38, 43, 98, 117, 119, 120, 143, 163, 169 |
| Larval body size | 1 | 2 | 66 |
| Leaf length | 1 | 3 | 20 |
| Leaf mass per unit area | 1 | 1 | 113 |
| Leaf water content | 1 | 1 | 113 |
| Length of corolla tube | 1 | 4 | 159 |
| Location | 39 | 156 | 3, 7, 8, 16, 25, 26, 38, 42, 44, 50, 61, 67, 70, 72, 77, 78, 82, 84, 88, 89, 90, 94, 95, 100, 102, 104, 105, 116, 121, 127, 137, 142, 159, 162, 163, 166, 168, 170, 171 |
| Location change | 2 | 26 | 77, 120 |
| Longevity | 24 | 61 | 3, 20, 22, 34, 50, 51, 55, 61, 66, 71, 88, 94, 101, 113, 118, 125, 127, 132, 138, 139, 147, 150, 151, 172 |
| Male combat | 1 | 2 | 141 |
| Male horn length | 1 | 2 | 21 |
| Mass-specific metabolic rate | 3 | 12 | 38, 40, 68 |
| Microhabitat breadth | 1 | 4 | 76 |
| Microhabitat type | 36 | 302 | 4, 7, 8, 15, 17, 25, 26, 34, 46, 47, 54, 59, 69, 70, 72, 80, 82, 88, 91, 95, 96, 98, 108, 120, 127, 138, 142, 143, 147, 150, 152, 167, 169, 171, 172, 173 |
| Migration distance | 24 | 90 | 2, 4, 7, 36, 39, 50, 67, 71, 73, 77, 78, 86, 87, 90, 93, 102, 106, 127, 134, 153, 154, 162, 168, 173 |
| Monogamy | 7 | 12 | 2, 7, 11, 61, 103, 115, 139 |
| Mouth brooding | 1 | 1 | 11 |
| Nest guarding | 3 | 6 | 11, 88, 125 |
| Nest habitat | 5 | 23 | 61, 67, 137, 143, 162 |
| Nest microhabitat | 20 | 75 | 2, 4, 45, 56, 68, 69, 71, 72, 77, 78, 88, 98, 108, 127, 128, 142, 143, 157, 162, 168 |
| Nest vulnerability | 2 | 6 | 56, 173 |
| Niche uniqueness | 1 | 2 | 154 |
| Number of reproductive years | 1 | 17 | 120 |
| Number of teats | 1 | 8 | 64 |
| Number of teats variability | 1 | 8 | 64 |
| Overwintering stage | 11 | 34 | 9, 14, 46, 52, 76, 110, 111, 112, 120, 123, 137 |
| Parental care | 10 | 25 | 7, 29, 66, 72, 73, 88, 125, 127, 143, 168 |
| Plant coverage | 1 | 3 | 92 |
| Plant growth form | 13 | 46 | 20, 94, 97, 118, 120, 150, 151, 156, 159, 165, 166, 167, 171 |

| | | | |
|---|---|---|---|
| Ploidy | 1 | 2 | 151 |
| Ploidy variability | 1 | 3 | 151 |
| Plumage polymorphism | 1 | 2 | 90 |
| Pollen vector | 9 | 27 | 62, 91, 94, 118, 120, 147, 150, 151, 156 |
| Pollution tolerance | 1 | 2 | 120 |
| Population abundance | 6 | 29 | 52, 93, 102, 119, 162, 168 |
| Population density | 18 | 82 | 9, 25, 26, 27, 38, 40, 53, 64, 65, 67, 69, 90, 96, 135, 139, 140, 150, 153 |
| Population density variability | 1 | 13 | 64 |
| Population growth rate | 6 | 21 | 55, 74, 75, 83, 131, 132 |
| Population juvenile ratio | 1 | 2 | 141 |
| Population level sex ratio | 1 | 2 | 141 |
| Precipitation | 16 | 107 | 15, 16, 25, 26, 33, 35, 46, 49, 71, 80, 93, 95, 116, 153, 164, 169 |
| Precipitation range | 6 | 27 | 15, 103, 141, 163, 164, 169 |
| Predator type | 1 | 2 | 127 |
| Productivity within range | 6 | 32 | 16, 38, 39, 43, 90, 169 |
| Proportion of range protected | 2 | 10 | 37, 43 |
| Protection status | 1 | 2 | 93 |
| Rate of climatic niche evolution | 1 | 4 | 93 |
| Rate of diversification | 2 | 10 | 41, 136 |
| Rate of habitat niche evolution | 1 | 4 | 93 |
| Rate of trophic niche evolution | 1 | 4 | 93 |
| Reproduction type (sexual or asexual) | 11 | 46 | 34, 62, 91, 113, 120, 147, 159, 163, 167, 171, 172 |
| Reproductive area | 1 | 10 | 120 |
| Seasonality of reproduction | 2 | 5 | 72, 143 |
| Seed bank longevity | 3 | 9 | 120, 147, 172 |
| Seed size | 5 | 17 | 20, 118, 147, 151, 171 |
| Sex change | 1 | 1 | 11 |
| Sexual dimorphism | 10 | 37 | 21, 82, 86, 90, 103, 114, 115, 128, 141, 157 |
| Shelter use | 1 | 5 | 99 |
| Social group size | 12 | 85 | 25, 26, 27, 38, 39, 40, 64, 69, 81, 103, 139, 140 |
| Social group size variability | 1 | 8 | 64 |
| Sociality | 10 | 33 | 2, 8, 27, 40, 67, 86, 120, 127, 168, 173 |
| Spawning aggregation | 2 | 2 | 11, 104 |
| Specific leaf area | 1 | 3 | 92 |
| Spire index | 1 | 9 | 30 |
| Survivability | 2 | 2 | 13, 71 |
| Tail length | 1 | 2 | 90 |
| Taxon age | 5 | 17 | 41, 57, 79, 91, 107 |
| Taxon richness | 2 | 5 | 41, 79 |
| Taxonomic group | 16 | 38 | 11, 16, 25, 37, 38, 39, 43, 48, 52, 72, 88, 89, 120, 121, 140, 143 |
| Temperature | 27 | 165 | 15, 16, 17, 25, 26, 29, 33, 35, 46, 48, 50, 61, 66, 71, 77, 78, 87, 88, 89, 93, 138, 153, 162, 164, 167, 168, 169 |
| Temperature for breeding | 3 | 9 | 29, 88, 127 |

| | | | |
|---|---|---|---|
| Temperature range | 12 | 53 | 15, 46, 77, 78, 98, 103, 131, 141, 145, 163, 164, 169 |
| Threat breadth | 3 | 26 | 16, 102, 107 |
| Threat range | 9 | 77 | 25, 26, 49, 85, 102, 106, 117, 119, 164 |
| Threat type | 7 | 131 | 5, 15, 16, 34, 102, 106, 163 |
| Timing of migration | 1 | 2 | 168 |
| Timing of reproductive season | 9 | 40 | 83, 87, 88, 92, 120, 127, 143, 147, 168 |
| Tongue length | 1 | 1 | 5 |
| Torpor/hibernation | 3 | 54 | 67, 68, 99 |
| Vegetative resprouting capacity | 2 | 2 | 156, 165 |
| Water balance within range | 1 | 2 | 169 |
| Weaning age | 10 | 76 | 25, 26, 43, 53, 63, 64, 65, 69, 135, 139 |
| Weaning age variability | 1 | 13 | 64 |
| Wing shape | 3 | 13 | 60, 81, 148 |
| Wing size | 13 | 47 | 10, 14, 42, 46, 52, 60, 71, 81, 82, 89, 90, 148, 160 |
| Wingtip shape | 1 | 2 | 148 |
| Wingtip size | 1 | 4 | 148 |
| Winter leaf carrying | 1 | 7 | 120 |

References

1. ABELSON, E.S. (2016) Brain size is correlated with endangerment status in mammals. *Proceedings of the Royal Society B: Biological Sciences* **283**, 20152772.

2. AMANO, T. & YAMAURA, Y. (2007) Ecological and life-history traits related to range contractions among breeding birds in Japan. *Biological Conservation* **137**, 271–282.

3. ANDERSON, S.C., FARMER, R.G., FERRETTI, F., HOUDE, A.L.S. & HUTCHINGS, J.A. (2011) Correlates of Vertebrate Extinction Risk in Canada. *BioScience* **61**, 538–549.

4. ANGERMEIER, P.L. (1995) Ecological Attributes of Extinction-Prone Species: Loss of Freshwater Fishes of Virginia. *Conservation Biology* **9**, 143–158.

5. ARBETMAN, M.P., GLEISER, G., MORALES, C.L., WILLIAMS, P. & AIZEN, M.A. (2017) Global decline of bumblebees is phylogenetically structured and inversely related to species range size and pathogen incidence. *Proceedings of the Royal Society B: Biological Sciences* **284**, 20170204.

6. ARBUCKLE, K. (2016) Chemical antipredator defence is linked to higher extinction risk. *Royal Society Open Science* **3**, 160681.

7. BARSHEP, Y., ERNI, B., UNDERHILL, L.G. & ALTWEGG, R. (2017) Identifying ecological and life-history drivers of population dynamics of wetland birds in South Africa. *Global Ecology and Conservation* **12**, 96–107.

8. BARTOMEUS, I., ASCHER, J.S., GIBBS, J., DANFORTH, B.N., WAGNER, D.L., HEDTKE, S.M. & WINFREE, R. (2013) Historical changes in northeastern US bee pollinators related to shared ecological traits. *Proceedings of the National Academy of Sciences* **110**, 4656–4660.

9. BARTONOVA, A., BENES, J. & KONVICKA, M. (2014) Generalist-specialist continuum and life history traits of Central European butterflies (Lepidoptera) - are we missing a part of the picture? *European Journal of Entomology*.

10. BASSET, Y., BARRIOS, H., SEGAR, S., SRYGLEY, R.B., AIELLO, A., WARREN, A.D., DELGADO, F., CORONADO, J., LEZCANO, J., ARIZALA, S., RIVERA, M., PEREZ, F., BOBADILLA, R., LOPEZ, Y. & RAMIREZ, J.A. (2015) The Butterflies of Barro Colorado Island, Panama: Local Extinction since the 1930s. *PLoS ONE* **10**, e0136623.

11. BENDER, M. G., FLOETER, S. R., MAYER, F. P., VILA-NOVA, D. A., LONGO, G. O., HANAZAKI, N., CARVALHO-FILHO, A. & FERREIRA, C. E. L. (2013) Biological attributes and major threats as predictors of the vulnerability of species: a case study with Brazilian reef fishes. *Oryx* **47**, 259–265.

12. BENNETT, P.M. & OWENS, I.P. (1997) Variation in extinction risk among birds: chance or evolutionary predisposition? *Proceedings of the Royal Society of London B: Biological Sciences* **264**, 401–408.

13. BENSCOTER, A.M., REECE, J.S., NOSS, R.F., BRANDT, L.A., MAZZOTTI, F.J., ROMAÑACH, S.S. & WATLING, J.I. (2013) Threatened and Endangered Subspecies with Vulnerable

Ecological Traits Also Have High Susceptibility to Sea Level Rise and Habitat Fragmentation. *PLoS ONE* **8**, e70647.

14. BETZHOLTZ, P.-E., FRANZÉN, M. & FORSMAN, A. (2017) Colour pattern variation can inform about extinction risk in moths. *Animal Conservation* **20**, 72–79.

15. BLAND, L.M. (2017) Global correlates of extinction risk in freshwater crayfish. *Animal Conservation* **20**, 532–542.

16. BÖHM, M., WILLIAMS, R., BRAMHALL, H.R., MCMILLAN, K.M., DAVIDSON, A.D., GARCIA, A., BLAND, L.M., BIELBY, J. & COLLEN, B. (2016) Correlates of extinction risk in squamate reptiles: the relative importance of biology, geography, threat and range size. *Global Ecology and Biogeography* **25**, 391–405.

17. BONELLI, S., CERRATO, C., LOGLISCI, N. & BALLETTO, E. (2011) Population extinctions in the Italian diurnal lepidoptera: an analysis of possible causes. *Journal of Insect Conservation* **15**, 879–890.

18. BOTTS, E.A., ERASMUS, B.F.N. & ALEXANDER, G.J. (2013) Small range size and narrow niche breadth predict range contractions in South African frogs. *Global Ecology and Biogeography* **22**, 567–576.

19. BOYLES, J.G. & STORM, J.J. (2007) The Perils of Picky Eating: Dietary Breadth Is Related to Extinction Risk in Insectivorous Bats. *PLoS ONE* **2**, e672.

20. BRADSHAW, C.J.A., GIAM, X., TAN, H.T.W., BROOK, B.W. & SODHI, N.S. (2008) Threat or invasive status in legumes is related to opposite extremes of the same ecological and life-history attributes: Twin fates of legume species. *Journal of Ecology* **96**, 869–883.

21. BRO-JØRGENSEN, J. (2014) Will their armaments be their downfall? Large horn size increases extinction risk in bovids: Large horn size increases extinction risk in bovids. *Animal Conservation* **17**, 80–87.

22. BUBOVÁ, T., KULMA, M., VRABEC, V. & NOWICKI, P. (2016) Adult longevity and its relationship with conservation status in European butterflies. *Journal of Insect Conservation* **20**, 1021–1032.

23. CARDILLO, M. (2003) Biological determinants of extinction risk: why are smaller species less vulnerable? *Animal Conservation* **6**, 63–69.

24. CARDILLO, M. & BROMHAM, L. (2001) Body Size and Risk of Extinction in Australian Mammals. *Conservation Biology* **15**, 1435–1440.

25. CARDILLO, M., MACE, G.M., GITTLEMAN, J.L., JONES, K.E., BIELBY, J. & PURVIS, A. (2008) The predictability of extinction: biological and external correlates of decline in mammals. *Proceedings of the Royal Society of London B: Biological Sciences* **275**, 1441–1448.

26. CARDILLO, M., MACE, G.M., JONES, K.E., BIELBY, J., BININDA-EMONDS, O.R.P., SECHREST, W., ORME, C.D.L. & PURVIS, A. (2005) Multiple Causes of High Extinction Risk in Large Mammal Species. *Science* **309**, 1239–1241.

27. CARDILLO, M., PURVIS, A., SECHREST, W., GITTLEMAN, J.L., BIELBY, J. & MACE, G.M. (2004) Human Population Density and Extinction Risk in the World's Carnivores. *PLoS Biol* **2**, e197.

28. CASSEY, P. (2001) Determining variation in the success of New Zealand land birds. *Global Ecology & Biogeography*, 161–172.

29. CHESSMAN, B.C. (2013) Identifying species at risk from climate change: Traits predict the drought vulnerability of freshwater fishes. *Biological Conservation* **160**, 40–49.

30. CHIBA, S. & ROY, K. (2011) Selectivity of terrestrial gastropod extinctions on an oceanic archipelago and insights into the anthropogenic extinction process. *Proceedings of the National Academy of Sciences* **108**, 9496–9501.

31. CHISHOLM, R.A. & TAYLOR, R. (2010) Body size and extinction risk in Australian mammals: An information-theoretic approach: INFORMATION THEORY AND EXTINCTION RISK. *Austral Ecology* **35**, 616–623.

32. COLLEN, B., BYKOVA, E., LING, S., MILNER-GULLAND, E.J. & PURVIS, A. (2006) Extinction Risk: A Comparative Analysis of Central Asian Vertebrates. *Biodiversity & Conservation* **15**, 1859–1871.

33. COLLEN, B., MCRAE, L., DEINET, S., DE PALMA, A., CARRANZA, T., COOPER, N., LOH, J. & BAILLIE, J.E.M. (2011) Predicting how populations decline to extinction. *Philosophical Transactions of the Royal Society B: Biological Sciences* **366**, 2577–2586.

34. COMEROS-RAYNAL, M.T., POLIDORO, B.A., BROATCH, J., MANN, B.Q., GORMAN, C., BUXTON, C.D., GOODPASTER, A.M., IWATSUKI, Y., MACDONALD, T.C., POLLARD, D., RUSSELL, B. & CARPENTER, K.E. (2016) Key predictors of extinction risk in sea breams and porgies (Family: Sparidae). *Biological Conservation* **202**, 88–98.

35. COOPER, N., BIELBY, J., THOMAS, G.H. & PURVIS, A. (2008) Macroecology and extinction risk correlates of frogs. *Global Ecology and Biogeography* **17**, 211–221.

36. CRIMMINS, S.M., BOMA, P. & THOGMARTIN, W.E. (2015) Projected Risk of Population Declines for Native Fish Species in the Upper Mississippi River. *River Research and Applications* **31**, 135–142.

37. DARRAH, S.E., BLAND, L.M., BACHMAN, S.P., CLUBBE, C.P. & TRIAS-BLASI, A. (2017) Using coarse-scale species distribution data to predict extinction risk in plants. *Diversity and Distributions* **23**, 435–447.

38. DAVIDSON, A.D., BOYER, A.G., KIM, H., POMPA-MANSILLA, S., HAMILTON, M.J., COSTA, D.P., CEBALLOS, G. & BROWN, J.H. (2012) Drivers and hotspots of extinction risk in marine mammals. *Proceedings of the National Academy of Sciences* **109**, 3395–3400.

39. DAVIDSON, A.D., HAMILTON, M.J., BOYER, A.G., BROWN, J.H. & CEBALLOS, G. (2009) Multiple ecological pathways to extinction in mammals. *Proceedings of the National Academy of Sciences* **106**, 10702–10705.

40. DAVIDSON, A.D., SHOEMAKER, K.T., WEINSTEIN, B., COSTA, G.C., BROOKS, T.M., CEBALLOS, G., RADELOFF, V.C., RONDININI, C. & GRAHAM, C.H. (2017) Geography of current and future global mammal extinction risk. *PLoS ONE* **12**, e0186934.

41. DAVIES, T.J., SMITH, G.F., BELLSTEDT, D.U., BOATWRIGHT, J.S., BYTEBIER, B., COWLING, R.M., FOREST, F., HARMON, L.J., MUASYA, A.M., SCHRIRE, B.D., STEENKAMP, Y., VAN DER BANK, M. & SAVOLAINEN, V. (2011) Extinction Risk and Diversification Are Linked in a Plant Biodiversity Hotspot. *PLoS Biology* **9**, e1000620.

42. DESENDER, K., DEKONINCK, W., DUFRÊNE, M. & MAES, D. (2010) Changes in the distribution of carabid beetles in Belgium revisited: Have we halted the diversity loss? *Biological Conservation* **143**, 1549–1557.

43. DI MARCO, M., BUCHANAN, G.M., SZANTOI, Z., HOLMGREN, M., GROTTOLO MARASINI, G., GROSS, D., TRANQUILLI, S., BOITANI, L. & RONDININI, C. (2014) Drivers of extinction risk in African mammals: the interplay of distribution state, human pressure, conservation response and species biology. *Philosophical Transactions of the Royal Society B: Biological Sciences* **369**, 20130198.

44. DUNCAN, J.R. & LOCKWOOD, J.L. (2001) Extinction in a field of bullets: a search for causes in the decline of the world's freshwater fishes. *Biological Conservation* **102**, 97–105.

45. DUNCAN, R.P. & BLACKBURN, T.M. (2004) Extinction and endemism in the New Zealand avifauna. *Global Ecology and Biogeography* **13**, 509–517.

46. ESSENS, T., VAN LANGEVELDE, F., VOS, R.A., VAN SWAAY, C.A.M. & WALLISDEVRIES, M.F. (2017) Ecological determinants of butterfly vulnerability across the European continent. *Journal of Insect Conservation* **21**, 439–450.

47. FATTORINI, S. (2013) Species ecological preferences predict extinction risk in urban tenebrionid beetle guilds. *Animal Biology* **63**, 93–106.

48. FIELD, I.C., MEEKAN, M.G., BUCKWORTH, R.C. & BRADSHAW, C.J.A. (2009) Chapter 4 Susceptibility of Sharks, Rays and Chimaeras to Global Extinction. In *Advances in Marine Biology* pp. 275–363. Elsevier.

49. FISHER, D.O., BLOMBERG, S.P. & OWENS, I.P. (2003) Extrinsic versus intrinsic factors in the decline and extinction of Australian marsupials. *Proceedings of the Royal Society of London B: Biological Sciences* **270**, 1801–1808.

50. FLOUSEK, J., TELENSKÝ, T., HANZELKA, J. & REIF, J. (2015) Population Trends of Central European Montane Birds Provide Evidence for Adverse Impacts of Climate Change on High-Altitude Species. *PLoS ONE* **10**, e0139465.

51. FORERO-MEDINA, G., VIEIRA, M.V., GRELLE, C.E. DE V. & ALMEIDA, P.J. (2009) Body size and extinction risk in Brazilian carnivores. *Biota Neotropica* **9**, 45–49.

52. FORISTER, M.L., JAHNER, J.P., CASNER, K.L., WILSON, J.S. & SHAPIRO, A.M. (2011) The race is not to the swift: Long-term data reveal pervasive declines in California's low-elevation butterfly fauna. *Ecology* **92**, 2222–2235.

53. FRITZ, S.A., BININDA-EMONDS, O.R.P. & PURVIS, A. (2009) Geographical variation in predictors of mammalian extinction risk: big is bad, but only in the tropics. *Ecology Letters* **12**, 538–549.

54. GAGE, G.S., BROOKE, M. DE L., SYMONDS, M.R.E. & WEGE, D. (2004) Ecological correlates of the threat of extinction in Neotropical bird species. *Animal Conservation* **7**, 161–168.

55. GARCIA, V.B., LUCIFORA, L.O. & MYERS, R.A. (2008) The importance of habitat and life history to extinction risk in sharks, skates, rays and chimaeras. *Proceedings of the Royal Society B: Biological Sciences* **275**, 83–89.

56. GARNETT, S.T. & BROOK, B.W. (2007) Modelling to forestall extinction of Australian tropical birds. *Journal of Ornithology* **148**, 311–320.

57. GASTON, K.J. (1997) Evolutionary age and risk of extinction in the global avifauna. *Evolutionary Ecology* **11**, 557–565.

58. GASTON, K.J. & BLACKBURN, T.M. (1995) Birds, body size and the threat of extinction. *Phil. Trans. R. Soc. Lond. B* **347**, 205–212.

59. GIAM, X., NG, T.H., LOK, A.F.S.L. & NG, H.H. (2011) Local geographic range predicts freshwater fish extinctions in Singapore: Extinction correlates of tropical freshwater fish. *Journal of Applied Ecology* **48**, 356–363.

60. GIBB, H., HJÄLTÉN, J., BALL, J.P., PETTERSSON, R.B., LANDIN, J., ALVINI, O. & DANELL, K. (2006) Wing loading and habitat selection in forest beetles: Are red-listed species poorer dispersers or more habitat-specific than common congenerics? *Biological Conservation* **132**, 250–260.

61. GLASS, W.R., CORKUM, L.D. & MANDRAK, N.E. (2017) Living on the edge: Traits of freshwater fish species at risk in Canada. *Aquatic Conservation: Marine and Freshwater Ecosystems* **27**, 938–945.

62. GODEFROID, S. (2014) Do plant reproductive traits influence species susceptibility to decline? *Plant Ecology and Evolution* **147**, 154–164.

63. GONZÁLEZ-SUÁREZ, M., GÓMEZ, A. & REVILLA, E. (2013) Which intrinsic traits predict vulnerability to extinction depends on the actual threatening processes. *Ecosphere* **4**, 1–16.

64. GONZÁLEZ-SUÁREZ, M. & REVILLA, E. (2013) Variability in life-history and ecological traits is a buffer against extinction in mammals. *Ecology Letters* **16**, 242–251.

65. GONZALEZ-VOYER, A., GONZÁLEZ-SUÁREZ, M., VILÀ, C. & REVILLA, E. (2016) Larger brain size indirectly increases vulnerability to extinction in mammals: BRAIN SIZE AND VULNERABILITY TO EXTINCTION. *Evolution* **70**, 1364–1375.

66. GRENOUILLET, G. & COMTE, L. (2014) Illuminating geographical patterns in species' range shifts. *Global Change Biology* **20**, 3080–3091.

67. HAMMERSON, G.A., KLING, M., HARKNESS, M., ORMES, M. & YOUNG, B.E. (2017) Strong geographic and temporal patterns in conservation status of North American bats. *Biological Conservation* **212**, 144–152.

68. Hanna, E. & Cardillo, M. (2013) A comparison of current and reconstructed historic geographic range sizes as predictors of extinction risk in Australian mammals. *Biological Conservation* **158**, 196–204.

69. Hanna, E. & Cardillo, M. (2014) Clarifying the relationship between torpor and anthropogenic extinction risk in mammals: Torpor and extinction risk in mammals. *Journal of Zoology* **293**, 211–217.

70. Hero, J.-M., Williams, S.E. & Magnusson, W.E. (2005) Ecological traits of declining amphibians in upland areas of eastern Australia. *Journal of Zoology* **267**, 221–232.

71. Hof, A.R., Rodríguez-Castañeda, G., Allen, A.M., Jansson, R. & Nilsson, C. (2017) Vulnerability of Subarctic and Arctic breeding birds. *Ecological Applications* **27**, 219–234.

72. Howard, S.D. & Bickford, D.P. (2014) Amphibians over the edge: silent extinction risk of Data Deficient species. *Diversity and Distributions* **20**, 837–846.

73. Jager, H.I., Rose, K.A. & Vila-Gispert, A. (2008) Life history correlates and extinction risk of capital-breeding fishes. *Hydrobiologia* **602**, 15–25.

74. Jennings, S., Greenstreet, S.P.R. & Reynolds, J.D. (1999) Structural change in an exploited fish community: a consequence of differential fishing effects on species with contrasting life histories. *Journal of Animal Ecology* **68**, 617–627.

75. Jennings, S., Reynolds, J.D. & Mills, S.C. (1998) Life history correlates of responses to fisheries exploitation. *Proceedings of the Royal Society of London B: Biological Sciences* **265**, 333–339.

76. Jeppsson, T. & Forslund, P. (2014) Species' traits explain differences in Red list status and long-term population trends in longhorn beetles: Traits and extinction risk in longhorn beetles. *Animal Conservation* **17**, 332–341.

77. Jiguet, F., Gadot, A.-S., Julliard, R., Newson, S.E. & Couvet, D. (2007) Climate envelope, life history traits and the resilience of birds facing global change. *Global Change Biology* **13**, 1672–1684.

78. Jiguet, F., Gregory, R.D., Devictor, V., Green, R.E., Voříšek, P., Van Strien, A. & Couvet, D. (2010) Population trends of European common birds are predicted by characteristics of their climatic niche. *Global Change Biology* **16**, 497–505.

79. Johnson, C.N., Delean, S. & Balmford, A. (2002) Phylogeny and the selectivity of extinction in Australian marsupials. *Animal Conservation* **5**, 135–142.

80. Johnson, C.N. & Isaac, J.L. (2009) Body mass and extinction risk in Australian marsupials: The 'Critical Weight Range' revisited. *Austral Ecology* **34**, 35–40.

81. Jones, K.E., Purvis, A. & Gittleman, J.L. (2003) Biological correlates of extinction risk in bats. *The American Naturalist* **161**, 601–614.

82. Jones, M.J., Fielding, A. & Sullivan, M. (2006) Analysing Extinction Risk in Parrots using Decision Trees. *Biodiversity & Conservation* **15**, 1993–2007.

83. JUAN-JORDÁ, M.J., MOSQUEIRA, I., FREIRE, J. & DULVY, N.K. (2015) Population declines of tuna and relatives depend on their speed of life. *Proceedings of the Royal Society B: Biological Sciences* **282**, 20150322.

84. KAMILAR, J.M. & PACIULLI, L.M. (2008) Examining the extinction risk of specialized folivores: a comparative study of Colobine monkeys. *American Journal of Primatology* **70**, 816–827.

85. KEANE, A., BROOKE, M. D. L. & MCGOWAN, P.J.K. (2005) Correlates of extinction risk and hunting pressure in gamebirds (Galliformes). *Biological Conservation* **126**, 216–233.

86. KOH, L.P., SODHI, N.S. & BROOK, B.W. (2004) Ecological Correlates of Extinction Proneness in Tropical Butterflies: Extinction Correlates of Tropical Butterflies. *Conservation Biology* **18**, 1571–1578.

87. KOLEČEK, J., ALBRECHT, T. & REIF, J. (2014) Predictors of extinction risk of passerine birds in a Central European country: Extinction risk of passerines. *Animal Conservation* **17**, 498–506.

88. KOPF, R.K., SHAW, C. & HUMPHRIES, P. (2017) Trait-based prediction of extinction risk of small-bodied freshwater fishes: Predicting Extinction Risk. *Conservation Biology* **31**, 581–591.

89. KOTZE, D.J. & O'HARA, R.B. (2003) Species decline - but why? Explanations of carabid beetle (Coleoptera, Carabidae) declines in Europe. *Oecologia* **135**, 138–148.

90. KRÜGER, O. & RADFORD, A.N. (2008) Doomed to die? Predicting extinction risk in the true hawks Accipitridae. *Animal Conservation* **11**, 83–91.

91. LAANISTO, L., SAMMUL, M., KULL, T., MACEK, P. & HUTCHINGS, M.J. (2015) Trait-based analysis of decline in plant species ranges during the 20th century: a regional comparison between the UK and Estonia. *Global Change Biology* **21**, 2726–2738.

92. LAUTERBACH, D., RÖMERMANN, C., JELTSCH, F. & RISTOW, M. (2013) Factors driving plant rarity in dry grasslands on different spatial scales: a functional trait approach. *Biodiversity and Conservation* **22**, 2337–2352.

93. LAVERGNE, S., EVANS, M.E.K., BURFIELD, I.J., JIGUET, F. & THUILLER, W. (2012) Are species' responses to global change predicted by past niche evolution? *Philosophical Transactions of the Royal Society B: Biological Sciences* **368**, 20120091–20120091.

94. LAVERGNE, S., MOLINA, J. & DEBUSSCHE, M. (2006) Fingerprints of environmental change on the rare mediterranean flora: a 115-year study. *Global Change Biology* **12**, 1466–1478.

95. LAWES, M.J., FISHER, D.O., JOHNSON, C.N., BLOMBERG, S.P., FRANK, A.S., FRITZ, S.A., MCCALLUM, H., VANDERWAL, J., ABBOTT, B.N. & LEGGE, S. (2015) Correlates of recent declines of rodents in northern and southern Australia: habitat structure is critical. *PLoS One* **10**, e0130626.

96. LEACH, K., KELLY, R., CAMERON, A., MONTGOMERY, W.I. & REID, N. (2015) Expertly Validated Models and Phylogenetically-Controlled Analysis Suggests Responses to Climate Change Are Related to Species Traits in the Order Lagomorpha. *PLoS ONE* **10**, e0122267.

97. Leão, T.C.C., Fonseca, C.R., Peres, C.A. & Tabarelli, M. (2014) Predicting Extinction Risk of Brazilian Atlantic Forest Angiosperms: Neotropical Plant Extinction Risk. *Conservation Biology* **28**, 1349–1359.

98. Lee, T.M. & Jetz, W. (2010) Unravelling the structure of species extinction risk for predictive conservation science. *Proceedings of the Royal Society of London B: Biological Sciences*, rspb20101877.

99. Liow, L.H., Fortelius, M., Lintulaakso, K., Mannila, H. & Stenseth, N.C. (2009) Lower Extinction Risk in Sleep-or-Hide Mammals. *The American Naturalist* **173**, 264–272.

100. Lips, K.R., Reeve, J.D. & Witters, L.R. (2003) Ecological Traits Predicting Amphibian Population Declines in Central America. *Conservation Biology* **17**, 1078–1088.

101. Liu, C., Comte, L. & Olden, J.D. (2017) Heads you win, tails you lose: Life-history traits predict invasion and extinction risk of the world's freshwater fishes: TRAITS PREDICT GLOBAL FRESHWATER FISH INVASION AND EXTINCTION. *Aquatic Conservation: Marine and Freshwater Ecosystems* **27**, 773–779.

102. Long, P.R., Székely, T., Kershaw, M. & O'Connell, M. (2007) Ecological factors and human threats both drive wildfowl population declines. *Animal Conservation* **10**, 183–191.

103. Lootvoet, A.C., Philippon, J. & Bessa-Gomes, C. (2015) Behavioral Correlates of Primates Conservation Status: Intrinsic Vulnerability to Anthropogenic Threats. *PLoS ONE* **10**, e0135585.

104. Luiz, O.J., Woods, R.M., Madin, E.M.P. & Madin, J.S. (2016) Predicting IUCN Extinction Risk Categories for the World's Data Deficient Groupers (Teleostei: Epinephelidae). *Conservation Letters* **9**, 342–350.

105. Mace, G.M., Collen, B., Fuller, R.A. & Boakes, E.H. (2010) Population and geographic range dynamics: implications for conservation planning. *Philosophical Transactions of the Royal Society B: Biological Sciences* **365**, 3743–3751.

106. Machado, N. & Loyola, R.D. (2013) A Comprehensive Quantitative Assessment of Bird Extinction Risk in Brazil. *PLoS ONE* **8**, e72283.

107. Mankga, L.T. & Yessoufou, K. (2017) Factors driving the global decline of cycad diversity. *AoB Plants* **9**, plx022.

108. Matsuzaki, S.-I.S., Takamura, N., Arayama, K., Tominaga, A., Iwasaki, J. & Washitani, I. (2011) Potential impacts of non-native channel catfish on commercially important species in a Japanese lake, as inferred from long-term monitoring data. *Aquatic Conservation: Marine and Freshwater Ecosystems* **21**, 348–357.

109. Matthews, L.J., Arnold, C., Machanda, Z. & Nunn, C.L. (2011) Primate extinction risk and historical patterns of speciation and extinction in relation to body mass. *Proceedings of the Royal Society B: Biological Sciences* **278**, 1256–1263.

110. MATTILA, N., KAITALA, V., KOMONEN, A., PÄIVINEN, J. & KOTIAHO, J.S. (2011) Ecological correlates of distribution change and range shift in butterflies: Distribution decline in butterflies. *Insect Conservation and Diversity* **4**, 239–246.

111. MATTILA, N., KOTIAHO, J.S., KAITALA, V. & KOMONEN, A. (2008) The use of ecological traits in extinction risk assessments: A case study on geometrid moths. *Biological Conservation* **141**, 2322–2328.

112. MATTILA, N., KOTIAHO, J.S., KAITALA, V., KOMONEN, A. & PÄIVINEN, J. (2009) Interactions between Ecological Traits and Host Plant Type Explain Distribution Change in Noctuid Moths. *Conservation Biology* **23**, 703–709.

113. MONKS, A. & BURROWS, L. (2014) Are threatened plant species specialists, or just more vulnerable to disturbance? *Journal of Applied Ecology* **51**, 1228–1235.

114. MORROW, E.H. & FRICKE, C. (2004) Sexual selection and the risk of extinction in mammals. *Proceedings of the Royal Society B: Biological Sciences* **271**, 2395–2401.

115. MORROW, E.H. & PITCHER, T.E. (2003) Sexual selection and the risk of extinction in birds. *Proceedings of the Royal Society of London B: Biological Sciences* **270**, 1793–1799.

116. MURPHY, B.P. & DAVIES, H.F. (2014) There is a critical weight range for Australia's declining tropical mammals: Critical weight range for Australia's declining tropical mammals. *Global Ecology and Biogeography* **23**, 1058–1061.

117. MURRAY, B.R. & HOSE, G.C. (2005) Life-history and ecological correlates of decline and extinction in the endemic Australian frog fauna. *Austral Ecology* **30**, 564–571.

118. MURRAY, B.R., THRALL, P.H. & LEPSCHI, B.J. (2002) Relating species rarity to life history in plants of eastern Australia, 14.

119. MURRAY, K.A., ROSAUER, D., MCCALLUM, H. & SKERRATT, L.F. (2011) Integrating species traits with extrinsic threats: closing the gap between predicting and preventing species declines. *Proceedings of the Royal Society of London B: Biological Sciences*, rspb20101872.

120. MUSTERS, C.J.M., KALKMAN, V. & VAN STRIEN, A. (2013) Predicting rarity and decline in animals, plants, and mushrooms based on species attributes and indicator groups. *Ecology and Evolution*, 3401–3414.

121. NÁJERA, A. & SIMONETTI, J.A. (2012) Threatened birds of Guatemala: a random subset of the avifauna? *Bird Conservation International* **22**, 348–353.

122. NORRIS, K. & HARPER, N. (2004) Extinction processes in hot spots of avian biodiversity and the targeting of pre–emptive conservation action. *Proceedings of the Royal Society of London B: Biological Sciences* **271**, 123–130.

123. NYLIN, S. & BERGSTRÖM, A. (2009) Threat status in butterflies and its ecological correlates: how far can we generalize? *Biodiversity and Conservation* **18**, 3243–3267.

124. OLDEN, J.D., HOGAN, Z.S. & ZANDEN, M.J.V. (2007) Small fish, big fish, red fish, blue fish: size-biased extinction risk of the world's freshwater and marine fishes. *Global Ecology and Biogeography* **16**, 694–701.

125. OLDEN, J.D., POFF, N.L. & BESTGEN, K.R. (2008) Trait synergisms and the rarity, extirpation, and extinction risk of desert fishes. *Ecology* **89**, 847–856.

126. OWENS, I.P.F. & BENNETT, P.M. (2000) Ecological basis of extinction risk in birds: Habitat loss versus human persecution and introduced predators. *Proceedings of the National Academy of Sciences* **97**, 12144–12148.

127. PARENT, S. & SCHRIML, L.M. (1995) A model for the determination of fish species at risk based upon life-history traits and ecological data. *Canadian Journal of Fisheries and Aquatic Sciences* **52**, 1768–1781.

128. PARLATO, E.H., ARMSTRONG, D.P. & INNES, J.G. (2015) Traits influencing range contraction in New Zealand's endemic forest birds. *Oecologia* **179**, 319–328.

129. PAYNE, J.L., BUSH, A.M., HEIM, N.A., KNOPE, M.L. & MCCAULEY, D.J. (2016) Ecological selectivity of the emerging mass extinction in the oceans. *Science*, aaf2416.

130. PEÑARANDA, D.A. & SIMONETTI, J.A. (2015) Predicting and setting conservation priorities for Bolivian mammals based on biological correlates of the risk of decline: Biological Correlates of Decline Risk. *Conservation Biology* **29**, 834–843.

131. PINSKY, M.L. & BYLER, D. (2015) Fishing, fast growth and climate variability increase the risk of collapse. *Proceedings of the Royal Society B: Biological Sciences* **282**, 20151053.

132. PINSKY, M.L., JENSEN, O.P., RICARD, D. & PALUMBI, S.R. (2011) Unexpected patterns of fisheries collapse in the world's oceans. *Proceedings of the National Academy of Sciences* **108**, 8317–8322.

133. PLEGUEZUELOS, J.M., BRITO, J.C., FAHD, S., FERICHE, M., MATEO, J.A., MORENO-RUEDA, G., REQUES, R. & SANTOS, X. (2010) Setting conservation priorities for the Moroccan herpetofauna: the utility of regional red lists. *Oryx* **44**, 501–508.

134. POCOCK, M.J.O. (2011) Can traits predict species' vulnerability? A test with farmland passerines in two continents. *Proceedings of the Royal Society B: Biological Sciences* **278**, 1532–1538.

135. POLAINA, E., REVILLA, E. & GONZÁLEZ-SUÁREZ, M. (2016) Putting susceptibility on the map to improve conservation planning, an example with terrestrial mammals. *Diversity and Distributions* **22**, 881–892.

136. POLISHCHUK, L.V., POPADIN, K.Y., BARANOVA, M.A. & KONDRASHOV, A.S. (2015) A genetic component of extinction risk in mammals. *Oikos* **124**, 983–993.

137. POWNEY, G.D., CHAM, S.S.A., SMALLSHIRE, D. & ISAAC, N.J.B. (2015) Trait correlates of distribution trends in the Odonata of Britain and Ireland. *PeerJ* **3**, e1410.

138. POWNEY, G.D., RAPACCIUOLO, G., PRESTON, C.D., PURVIS, A. & ROY, D.B. (2014) A phylogenetically-informed trait-based analysis of range change in the vascular plant flora of Britain. *Biodiversity and Conservation* **23**, 171–185.

139. PRICE, S.A. & GITTLEMAN, J.L. (2007) Hunting to extinction: biology and regional economy influence extinction risk and the impact of hunting in artiodactyls. *Proceedings of the Royal Society B: Biological Sciences* **274**, 1845–1851.

140. PURVIS, A., GITTLEMAN, J.L., COWLISHAW, G. & MACE, G.M. (2000) Predicting extinction risk in declining species. *Proceedings of the Royal Society of London B: Biological Sciences* **267**, 1947–1952.

141. REED, R.N. & SHINE, R. (2002) Lying in Wait for Extinction: Ecological Correlates of Conservation Status among Australian Elapid Snakes. *Conservation Biology* **16**, 451–461.

142. REYNOLDS, J.D., WEBB, T.J. & HAWKINS, L.A. (2005) Life history and ecological correlates of extinction risk in European freshwater fishes. *Canadian Journal of Fisheries and Aquatic Sciences* **62**, 854–862.

143. RIBEIRO, J., COLLI, G.R., CALDWELL, J.P. & SOARES, A.M.V.M. (2016) An integrated trait-based framework to predict extinction risk and guide conservation planning in biodiversity hotspots. *Biological Conservation* **195**, 214–223.

144. RIPPLE, W.J., WOLF, C., NEWSOME, T.M., HOFFMANN, M., WIRSING, A.J. & MCCAULEY, D.J. (2017) Extinction risk is most acute for the world's largest and smallest vertebrates. *Proceedings of the National Academy of Sciences*, 201702078.

145. ROLLAND, J. & SALAMIN, N. (2016) Niche width impacts vertebrate diversification. *Global Ecology and Biogeography* **25**, 1252–1263.

146. RULAND, F. & JESCHKE, J.M. (2017) Threat-dependent traits of endangered frogs. *Biological Conservation* **206**, 310–313.

147. SAAR, L., TAKKIS, K., PÄRTEL, M. & HELM, A. (2012) Which plant traits predict species loss in calcareous grasslands with extinction debt? *Diversity and Distributions* **18**, 808–817.

148. SAFI, K. & KERTH, G. (2004) A Comparative Analysis of Specialization and Extinction Risk in Temperate-Zone Bats. *Conservation Biology* **18**, 1293–1303.

149. SAGOT, M. & CHAVERRI, G. (2015) Effects of roost specialization on extinction risk in bats. *Conservation Biology* **29**, 1666–1673.

150. SAKAI, A.K., WAGNER, W.L. & MEHRHOFF, L.A. (2002) Patterns of Endangerment in the Hawaiian Flora. *Systematic Biology* **51**, 276–302.

151. SCHMIDT, J.P., STEPHENS, P.R. & DRAKE, J.M. (2012) Two sides of the same coin? Rare and pest plants native to the United States and Canada. *Ecological Applications* **22**, 1512–1525.

152. SEIBOLD, S., BRANDL, R., BUSE, J., HOTHORN, T., SCHMIDL, J., THORN, S. & MÜLLER, J. (2015) Association of extinction risk of saproxylic beetles with ecological degradation of

forests in Europe: Beetle Extinction and Forest Degradation. *Conservation Biology* **29**, 382–390.

153. SEOANE, J. & CARRASCAL, L.M. (2007) Interspecific differences in population trends of Spanish birds are related to habitat and climatic preferences. *Global Ecology and Biogeography* **17**, 111–121.

154. SHULTZ, S., B. BRADBURY, R., L. EVANS, K., D. GREGORY, R. & M. BLACKBURN, T. (2005) Brain size and resource specialization predict long-term population trends in British birds. *Proceedings of the Royal Society B: Biological Sciences* **272**, 2305–2311.

155. SILICEO, I. & DÍAZ, J.A. (2010) A comparative study of clutch size, range size, and the conservation status of island vs. mainland lacertid lizards. *Biological Conservation* **143**, 2601–2608.

156. SODHI, N.S., KOH, L.P., PEH, K.S.-H., TAN, H.T.W., CHAZDON, R.L., CORLETT, R.T., LEE, T.M., COLWELL, R.K., BROOK, B.W., SEKERCIOGLU, C.H. & BRADSHAW, C.J.A. (2008) Correlates of extinction proneness in tropical angiosperms. *Diversity and Distributions* **14**, 1–10.

157. SODHI, N.S., LEE, T.M., KOH, L.P. & PRAWIRADILAGA, D.M. (2006) Long-Term Avifaunal Impoverishment in an Isolated Tropical Woodlot. *Conservation Biology* **20**, 772–779.

158. SREEKAR, R., HUANG, G., ZHAO, J.-B., PASION, B.O., YASUDA, M., ZHANG, K., PEABOTUWAGE, I., WANG, X., QUAN, R.-C., FERRY SLIK, J.W., CORLETT, R.T., GOODALE, E. & HARRISON, R.D. (2015) The use of species-area relationships to partition the effects of hunting and deforestation on bird extirpations in a fragmented landscape. *Diversity and Distributions* **21**, 441–450.

159. STEFANAKI, A., KANTSA, A., TSCHEULIN, T., CHARITONIDOU, M. & PETANIDOU, T. (2015) Lessons from Red Data Books: Plant Vulnerability Increases with Floral Complexity. *PLoS ONE* **10**, e0138414.

160. SULLIVAN, M.S., GILBERT, F., ROTHERAY, G., CROASDALE, S. & JONES, M. (2000) Comparative analyses of correlates of Red data book status: a case study using European hoverflies (Diptera: Syrphidae). *Animal Conservation* **3**, 91–95.

161. TERZOPOULOU, S., RIGAL, F., WHITTAKER, R.J., BORGES, P.A.V. & TRIANTIS, K.A. (2015) Drivers of extinction: the case of Azorean beetles. *Biology Letters* **11**, 20150273.

162. THAXTER, C.B., JOYS, A.C., GREGORY, R.D., BAILLIE, S.R. & NOBLE, D.G. (2010) Hypotheses to explain patterns of population change among breeding bird species in England. *Biological Conservation* **143**, 2006–2019.

163. TINGLEY, R., HITCHMOUGH, R.A. & CHAPPLE, D.G. (2013) Life-history traits and extrinsic threats determine extinction risk in New Zealand lizards. *Biological Conservation* **165**, 62–68.

164. TINGLEY, R., MAHONEY, P.J., DURSO, A.M., TALLIAN, A.G., MORÁN-ORDÓÑEZ, A. & BEARD, K.H. (2016) Threatened and invasive reptiles are not two sides of the same coin: Extinction and invasion risk in reptiles. *Global Ecology and Biogeography* **25**, 1050–1060.

165. TRINDER-SMITH, H., COWLING, R.M. & LINDER, H.P. (1996) Profiling a besieged flora: endemic and threatened plants of the Cape Peninsula, South Africa. *Biodiversity and Conservation* **5**, 575–589.

166. VAMOSI, J.C. & VAMOSI, S.M. (2005) Present day risk of extinction may exacerbate the lower species richness of dioecious clades: Extinction risk of dioecious plants. *Diversity and Distributions* **11**, 25–32.

167. VAN CALSTER, H., VANDENBERGHE, R., RUYSEN, M., VERHEYEN, K., HERMY, M. & DECOCQ, G. (2008) Unexpectedly high 20th century floristic losses in a rural landscape in northern France: Floristic changes in rural landscapes. *Journal of Ecology* **96**, 927–936.

168. VAN TURNHOUT, C.A.M., FOPPEN, R.P.B., LEUVEN, R.S.E.W., VAN STRIEN, A. & SIEPEL, H. (2010) Life-history and ecological correlates of population change in Dutch breeding birds. *Biological Conservation* **143**, 173–181.

169. VERDE ARREGOITIA, L.D., LEACH, K., REID, N. & FISHER, D.O. (2015) Diversity, extinction, and threat status in Lagomorphs. *Ecography* **38**, 1155–1165.

170. VILCHIS, L.I., JOHNSON, C.K., EVENSON, J.R., PEARSON, S.F., BARRY, K.L., DAVIDSON, P., RAPHAEL, M.G. & GAYDOS, J.K. (2015) Assessing ecological correlates of marine bird declines to inform marine conservation: Ecological Correlates of Seabird Declines. *Conservation Biology* **29**, 154–163.

171. WALKER, K.J. & PRESTON, C.D. (2006) Ecological Predictors of Extinction Risk in the Flora of Lowland England, UK. *Biodiversity & Conservation* **15**, 1913–1942.

172. WALKER, K.J., PRESTON, C.D. & BOON, C.R. (2009) Fifty years of change in an area of intensive agriculture: plant trait responses to habitat modification and conservation, Bedfordshire, England. *Biodiversity and Conservation* **18**, 3597–3613.

173. WANG, Y., SI, X., BENNETT, P.M., CHEN, C., ZENG, D., ZHAO, Y., WU, Y. & DING, P. (2018) Ecological correlates of extinction risk in Chinese birds. *Ecography* **41**, 782–794.

#### Appendix S2 – Tables with all information on studies and traits collected.

The information is stored in an excel file ("Chichorro et al__SuppInfo_S2.xls"). Three tables are available in separate sheets (studies, tests, variables) as well as a legend.

Appendix S3 – Proportion of significant measurements by trait and taxon

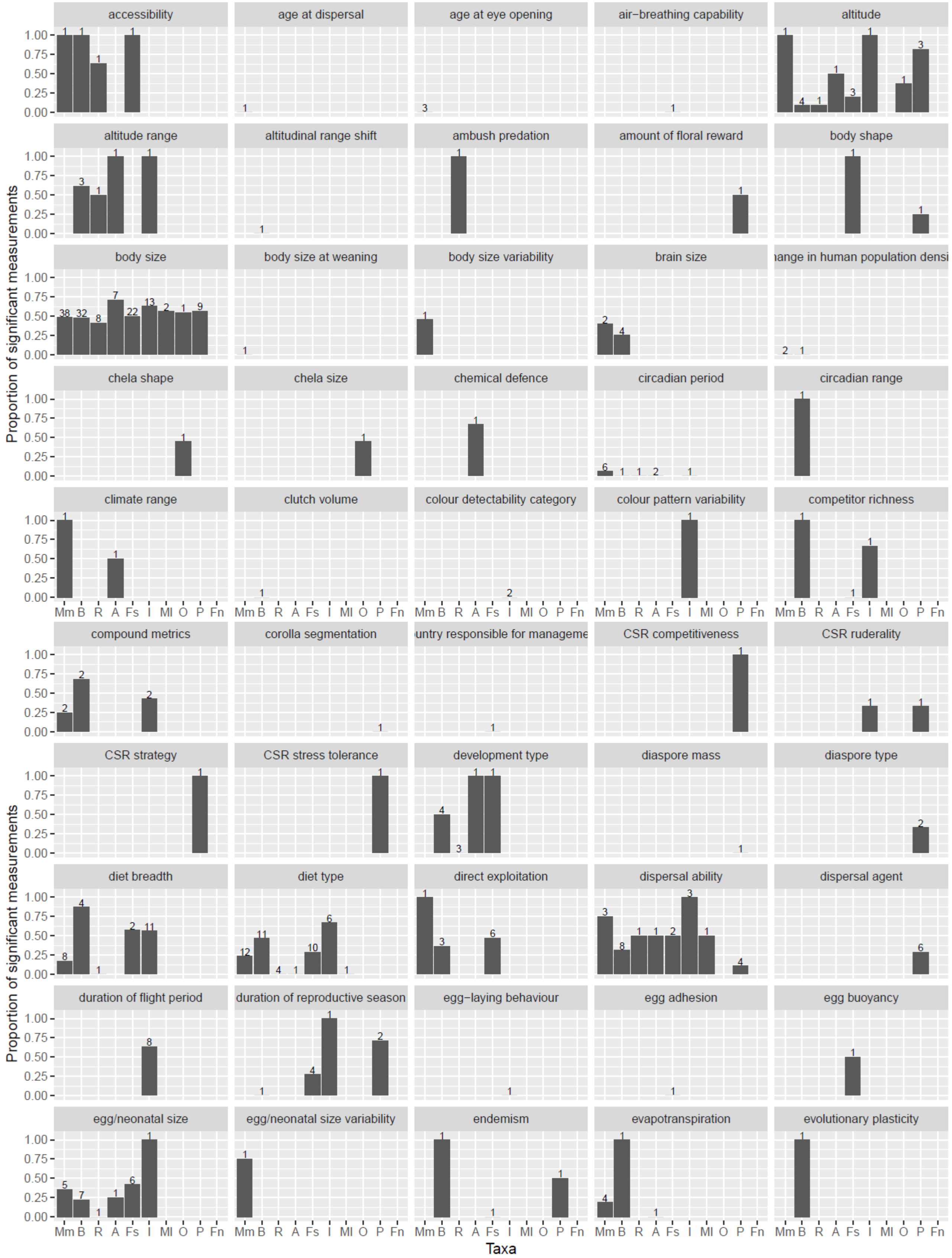


Figure S1: Proportion of significant measurements per trait (part 2, traits are ordered alphabetically), per taxon. The number of studies is given on top of each bar. **Mm:** Mammals, **B**: Birds, **R**: Reptiles, **A**: Amphibians, **Fs**: Fishes, **I**: Insects, **Ml**: Molluscs, **O**: Other Invertebrates, **P**: Plants, **Fn**: Fungi.

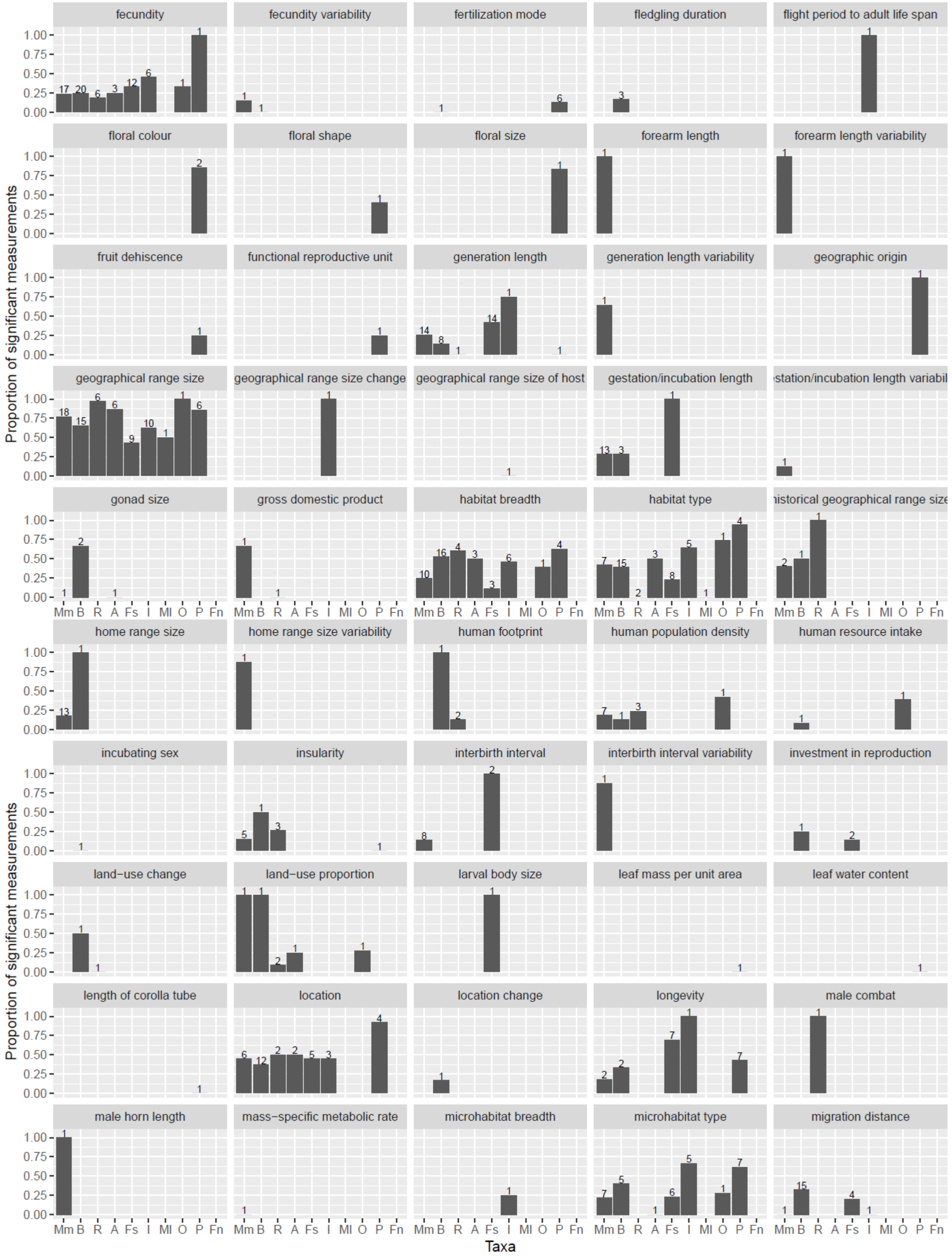


Figure S2: Proportion of significant measurements per trait (part 2, traits are ordered alphabetically), per taxon. The number of studies is given on top of each bar. **Mm:** Mammals, **B**: Birds, **R**: Reptiles, **A**: Amphibians, **Fs**: Fishes, **I**: Insects, **Ml**: Molluscs, **O**: Other Invertebrates, **P**: Plants, **Fn**: Fungi.

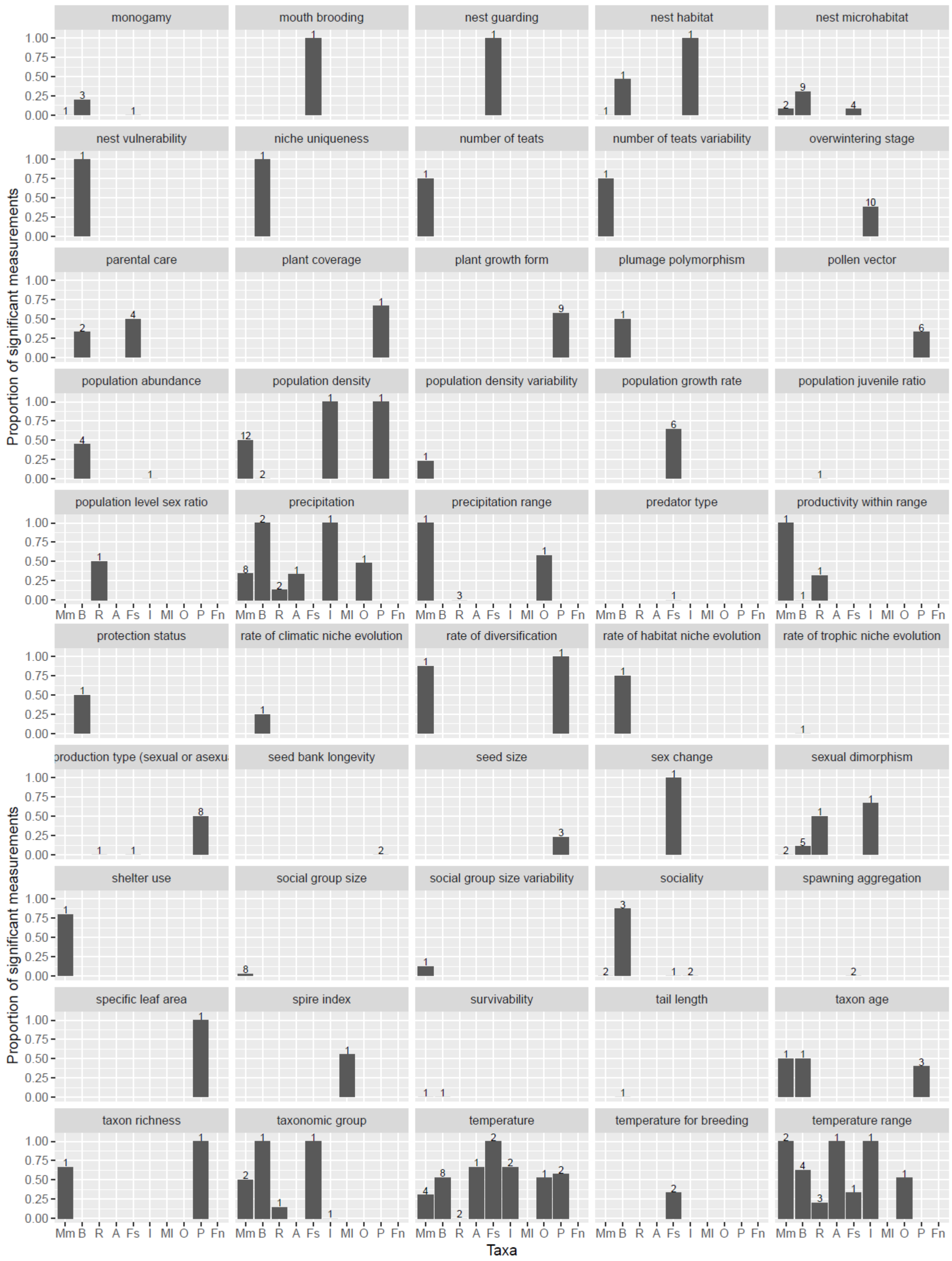


Figure S3: Proportion of significant measurements per trait (part 3, traits are ordered alphabetically), per taxon. The number of studies is given on top of each bar. **Mm:** Mammals, **B**: Birds, **R**: Reptiles, **A**: Amphibians, **Fs**: Fishes, **I**: Insects, **Ml**: Molluscs, **O**: Other Invertebrates, **P**: Plants, **Fn**: Fungi.

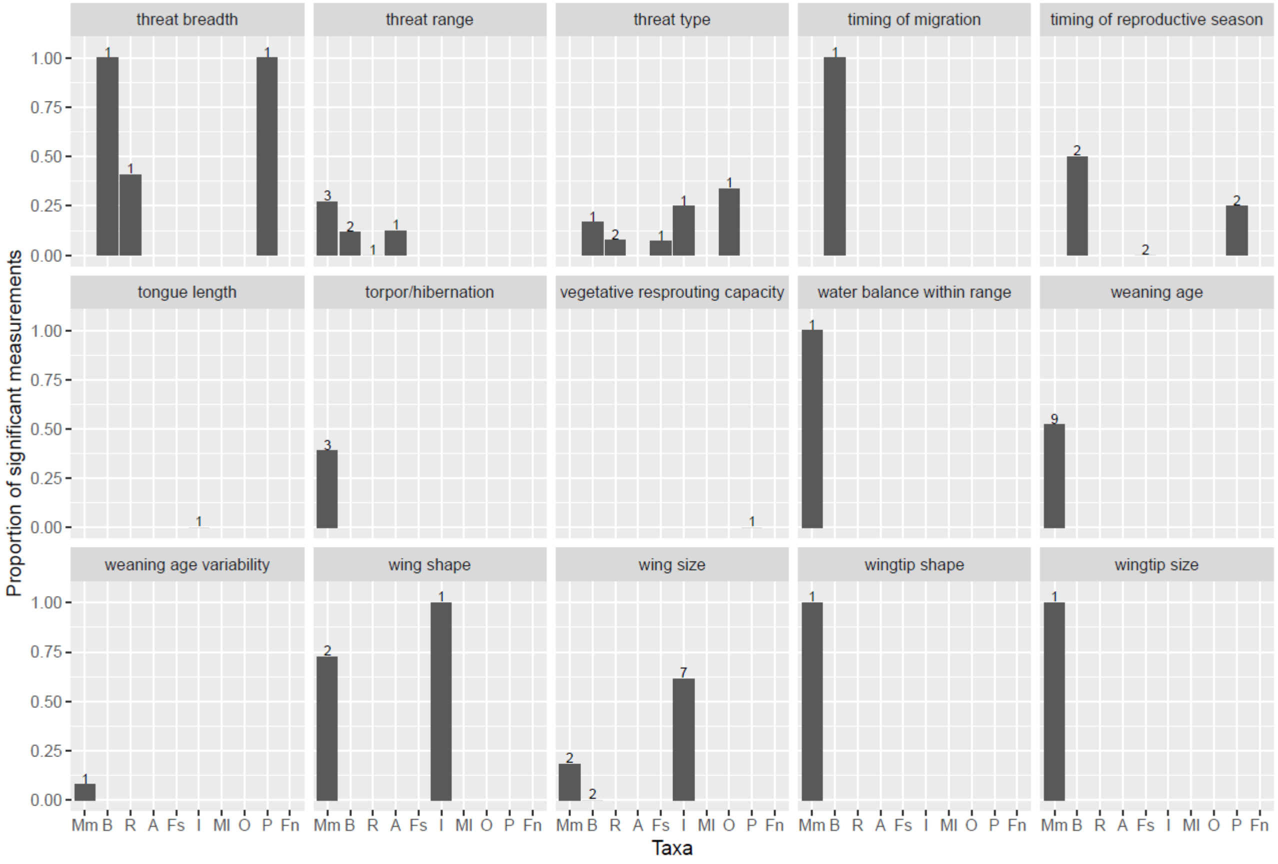


Figure S4: Proportion of significant measurements per trait (part 4, traits are ordered alphabetically), per taxon. The number of studies is given on top of each bar. **Mm:** Mammals, **B**: Birds, **R**: Reptiles, **A**: Amphibians, **Fs**: Fishes, **I**: Insects, **Ml**: Molluscs, **O**: Other Invertebrates, **P**: Plants, **Fn**: Fungi.

Appendix S4 – Forest plots of the meta-analyses

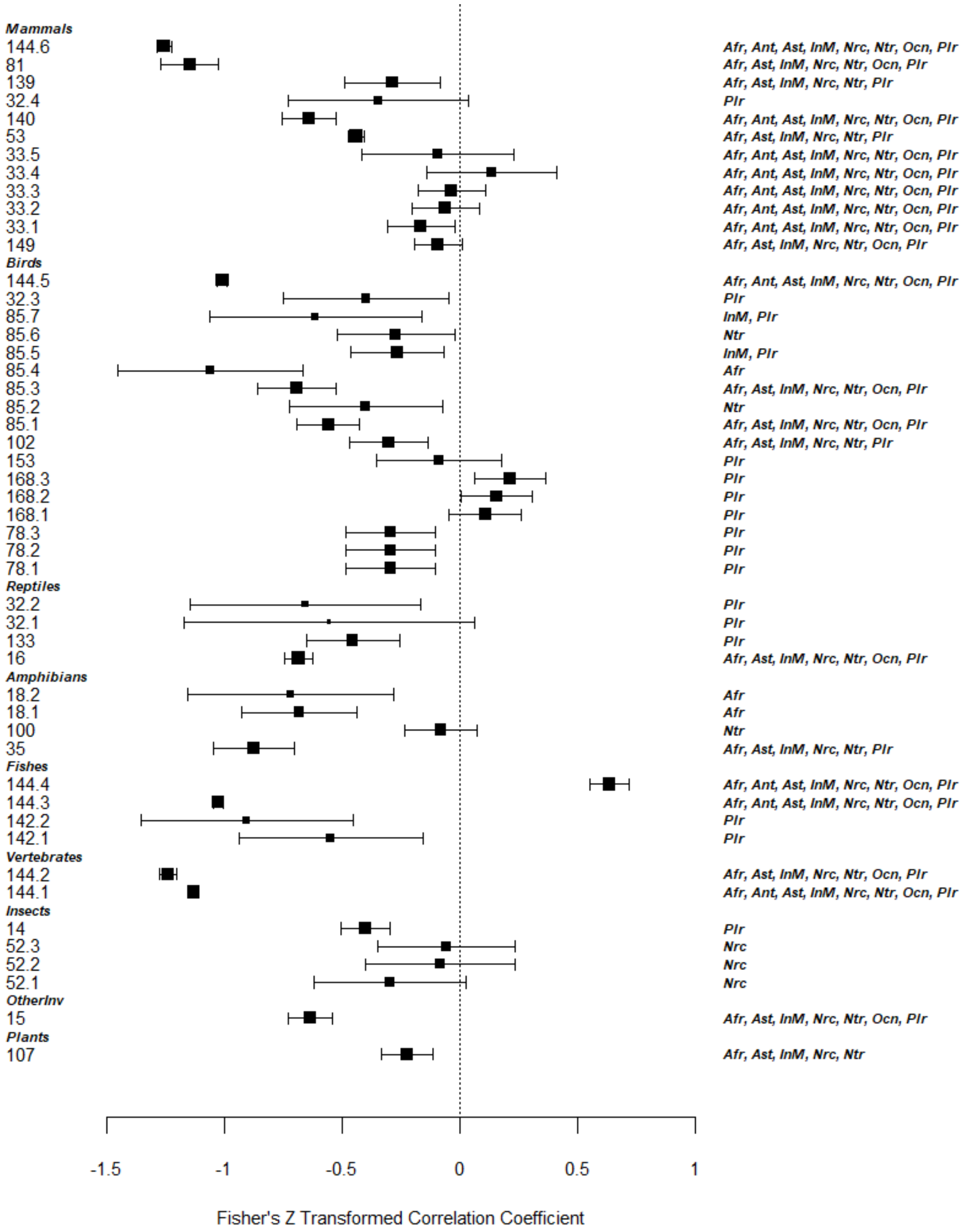


Figure S5: Forest plot of a random-effects model of the effect of geographical range size on extinction risk. Higher values of Fisher's Z correlation coefficients indicate higher extinction risk associated with higher values of geographical range size. In the left-hand labels, the identity of each effect size is shown in the format '<Study Identifier from Appendix S1>.<measurement identifier>'. To the right are the biogeographical realms included in each of the tests, where **Afr:** Afrotropics, **Ant:** Antarctic, **Ast:** Australasian, **InM:** Indo-Malaya, **Nrc:** Nearctic, **Ntr:** Neotropics, **Ocn:** Oceania, **Plr:** Palaearctic.

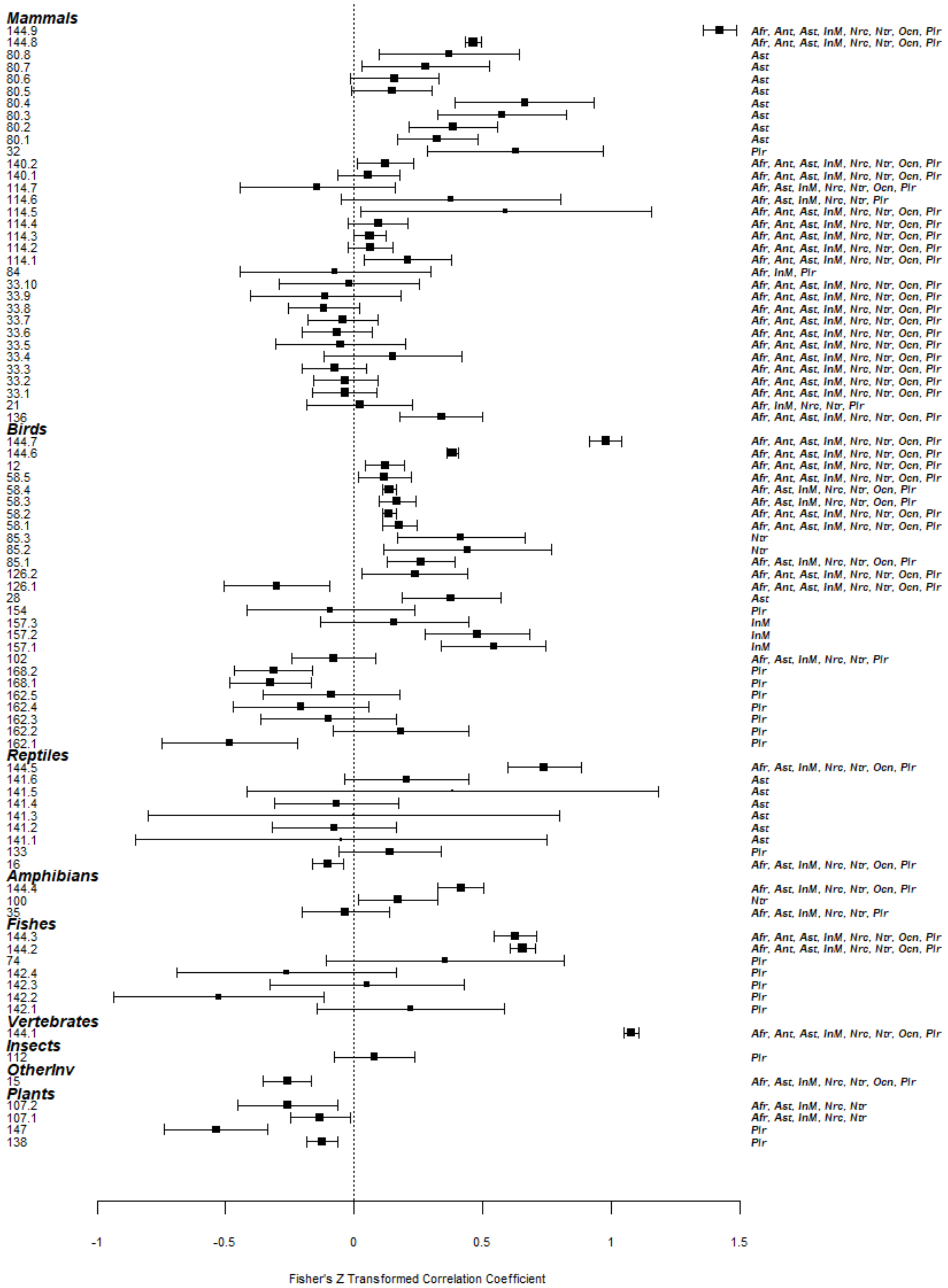


Figure S6: Forest plot of a random-effects model of the effect of body size on extinction risk. Higher values of Fisher's Z correlation coefficients indicate higher extinction risk associated with higher values of body size. In the left-hand labels, the identity of each effect size is shown in the format '<Study Identifier from Appendix S1>.<measurement identifier>'. To the right are the biogeographical realms included in each of the tests, where **Afr:** Afrotropics, **Ant:** Antarctic, **Ast:** Australasian, **InM:** Indo-Malaya, **Nrc:** Nearctic, **Ntr:** Neotropics, **Ocn:** Oceania, **Plr:** Palaearctic.

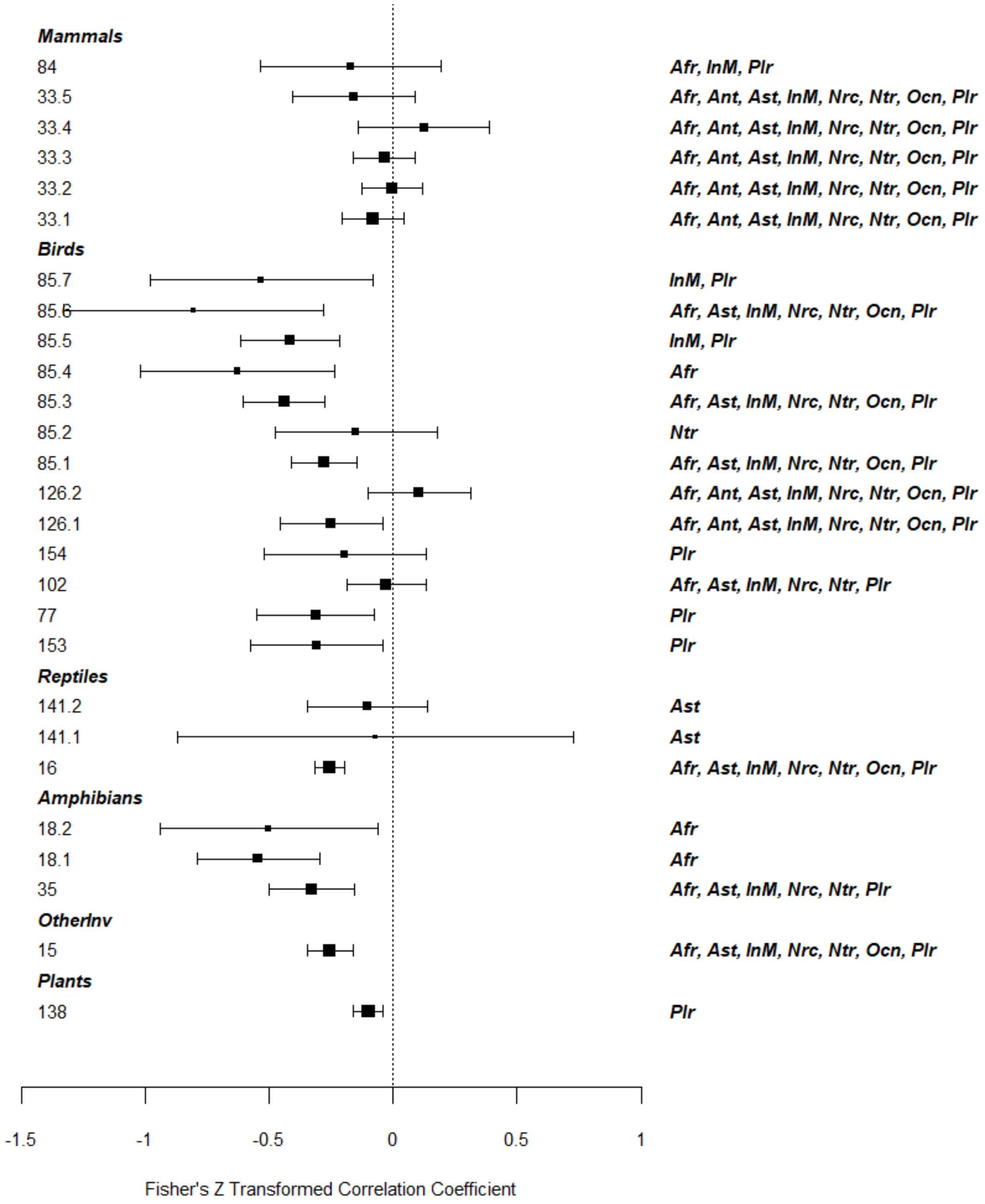


Figure S7: Forest plot of a random-effects model of the effect of habitat breadth on extinction risk. Higher values of Fisher's Z correlation coefficients indicate higher extinction risk associated with higher values of habitat breadth. In the left-hand labels, the identity of each effect size is shown in the format '<Study Identifier from Appendix S1>.<measurement identifier>'. To the right are the biogeographical realms included in each of the tests, where **Afr:** Afrotropics, **Ant:** Antarctic, **Ast:** Australasian, **InM:** Indo-Malaya, **Nrc:** Nearctic, **Ntr:** Neotropics, **Ocn:** Oceania, **Plr:** Palaearctic.
